# Rapid activation of IL-2 receptor signaling by CD301b^+^ DC-derived IL-2 dictates the outcome of helper T cell differentiation

**DOI:** 10.1101/2023.10.26.564276

**Authors:** Naoya Tatsumi, Jihad El-Fenej, Alejandro Davila-Pagan, Yosuke Kumamoto

## Abstract

Effector T helper (Th) cell differentiation is fundamental to functional adaptive immunity. Different subsets of dendritic cells (DCs) preferentially induce different types of Th cells, but the fate instruction mechanism for Th type 2 (Th2) differentiation remains enigmatic, as the critical DC-derived cue has not been clearly identified. Here, we show that CD301b^+^ DCs, a major Th2-inducing DC subset, drive Th2 differentiation through cognate interaction by ‘kick-starting’ IL-2 receptor signaling in CD4T cells. Mechanistically, CD40 engagement induces IL-2 production selectively from CD301b^+^ DCs to maximize CD25 expression in CD4 T cells, which is required specifically for the Th2 fate decision. On the other hand, CD25 in CD301b^+^ DCs facilitates directed action of IL-2 toward cognate CD4T cells. Furthermore, CD301b^+^ DC-derived IL-2 skews CD4T cells away from the T follicular helper fate. These results highlight the critical role of DC-intrinsic CD40–IL-2 axis in bifurcation of Th cell fate.

## Introduction

Activation of antigen-specific CD4T cells by dendritic cells (DCs) is critical for initiating adaptive immune responses. Upon recognition of cognate peptide-major histocompatibility complex class II (MHCII) complex through the T cell receptor (TCR), naive CD4T cells rapidly proliferate as they differentiate into distinct subsets of effector T helper (Th) cells such as Th1, Th2, and Th17 cells, each of which have unique functions in immunity ^1^. DCs play a crucial role in both expansion and differentiation of antigen-specific CD4T cells not only by presenting antigens with costimulation but also by producing cytokines that affect the Th cell fate decision ^2^. Notably, different DC subsets have been shown to produce distinct patterns of cytokines upon stimulation and thereby preferentially induce different types of Th cells. For instance, among the three major DC subsets in mouse skin, including the dermal type 1 conventional DCs (cDC1s), the dermal cDC2s, and the epidermal Langerhans cells (LCs), which have a monocytic origin but have DC-like function, the dermal cDC1s preferentially induce Th1 cell differentiation by producing IL-12 ^3, 4^, whereas LCs induce Th17 cell differentiation through IL-6 production ^3, 5^. Such division of labor between DC subsets is not unique to the skin, although, in organs outside of the skin where LCs do not exist, the Th17 cell differentiation is induced by a specialized subset of cDC2s that produce high levels of IL-23 and/or IL-6 ^6, 7, 8, 9^. In contrast, the dermal cDC2s and cDC2s elsewhere have been shown to be required for the differentiation of Th2 cells ^10, 11, 12, 13, 14^, which play a crucial role in the development of allergic diseases and protection against helminth parasites. However, unlike the Th1 and Th17 fate induced by defined cytokines derived from a particular DC subset, the DC-derived factors acting directly on CD4T cells for instructing the Th2 cell fate have not been clearly identified.

We and others have shown previously that CD301b^+^ DCs, a major, migratory subset of cDC2s, are required specifically for the Th2 cell differentiation of antigen-specific CD4T cells induced by allergens, adjuvants, or by helminth infection ^10, 11, 15, 16^. Diphtheria toxin (DT)-induced depletion of CD301b^+^ DCs in mice expressing the DT receptor (DTR) under the regulation of the *Mgl2* (encoding CD301b) promoter (Mgl2-DTR mice) resulted not only in delayed priming but also in reduced Th2 cell differentiation with a compensatory increase in Th1, Th17 and T follicular helper (Tfh) cell differentiation ^17, 18^. These findings suggest that CD301b^+^ DCs dictate the bifurcation of effector CD4T cell fate, but its mechanism remains unknown. Here, we show that the Th2 cell fate instruction by CD301b^+^ DCs requires MHCII– and CD40-dependent cognate interaction with CD4T cells. Mechanistically, the CD40 ligation induces IL-2 production specifically from CD301b^+^ DCs, which is required for the maximal CD25 upregulation and the downstream IL-2 receptor (IL-2R) signaling in antigen-specific CD4T cells. The full expression of CD25 in antigen-specific CD4T cells is required for their differentiation into Th2 but not Th1 cells. In addition, the IL-2R signaling in the cognate CD4T cells early after priming requires CD25 expression in CD301b^+^ DCs, suggesting that the DC-derived CD25 facilitates the directed action of IL-2. Furthermore, the CD301b^+^ DC-intrinsic CD40–IL2 axis skews CD4T cells away from differentiation into Tfh cells. These data highlight a novel role for DC-intrinsic CD40–IL-2 axis in bifurcation of Th cell fate.

## Results

### Direct antigen presentation by CD301b^+^ DCs is required for Th2 cell differentiation

CD301b^+^ DCs are required for Th2 cell differentiation induced by papain (protease allergen), alum (type 2 adjuvant), and *Nippostrongylus brasiliensis* (helminth parasite) ^11^, but whether they instruct the Th2 cell fate through a cognate interaction or via antigen-independent cues (e.g., cytokines) remains unclear. To address this question, we examined Th2 cell differentiation in mice lacking MHCII specifically in CD301b^+^ DCs (*Mgl2^+/Cre^*;*H2ab1^fl/fl^*mice, hereafter CD301b^ΔMHCII^ mice) ^18^ by adoptively transferring carboxyfluorescein succinimidyl ester (CFSE)-labeled ovalbumin (OVA)-specific TCR transgenic CD4T (OT-II) cells ^19^ and subcutaneously immunizing in the footpad with OVA plus papain (**Fig. 1A**). Seven days after the immunization, the OT-II cells in the draining lymph node (dLN) of the CD301b^ΔMHCII^ mice showed normal cell divisions with a mild increase in number (**Fig. 1B,C**), indicating that they were primed by antigen-presenting cells other than CD301b^+^ DCs. However, as in CD301b^+^ DC-depleted mice ^11^, the OT-II cells in CD301b^ΔMHCII^ mice failed to develop IL-4-producing Th2 cells, while their differentiation into IFNγ-producing Th1 cells remained unchanged (**Fig. 1D,E**). Likewise, when mice were immunized with OVA plus alum, the Th2 differentiation of OT-II cells was impaired in CD301b^ΔMHCII^ mice without affecting cell divisions, numbers, or differentiation into Th1 cells (**Fig. 1F-H**). These results indicate that cognate interaction with CD301b^+^ DC is required for the differentiation of Th2 cells.

**Fig. 1.**
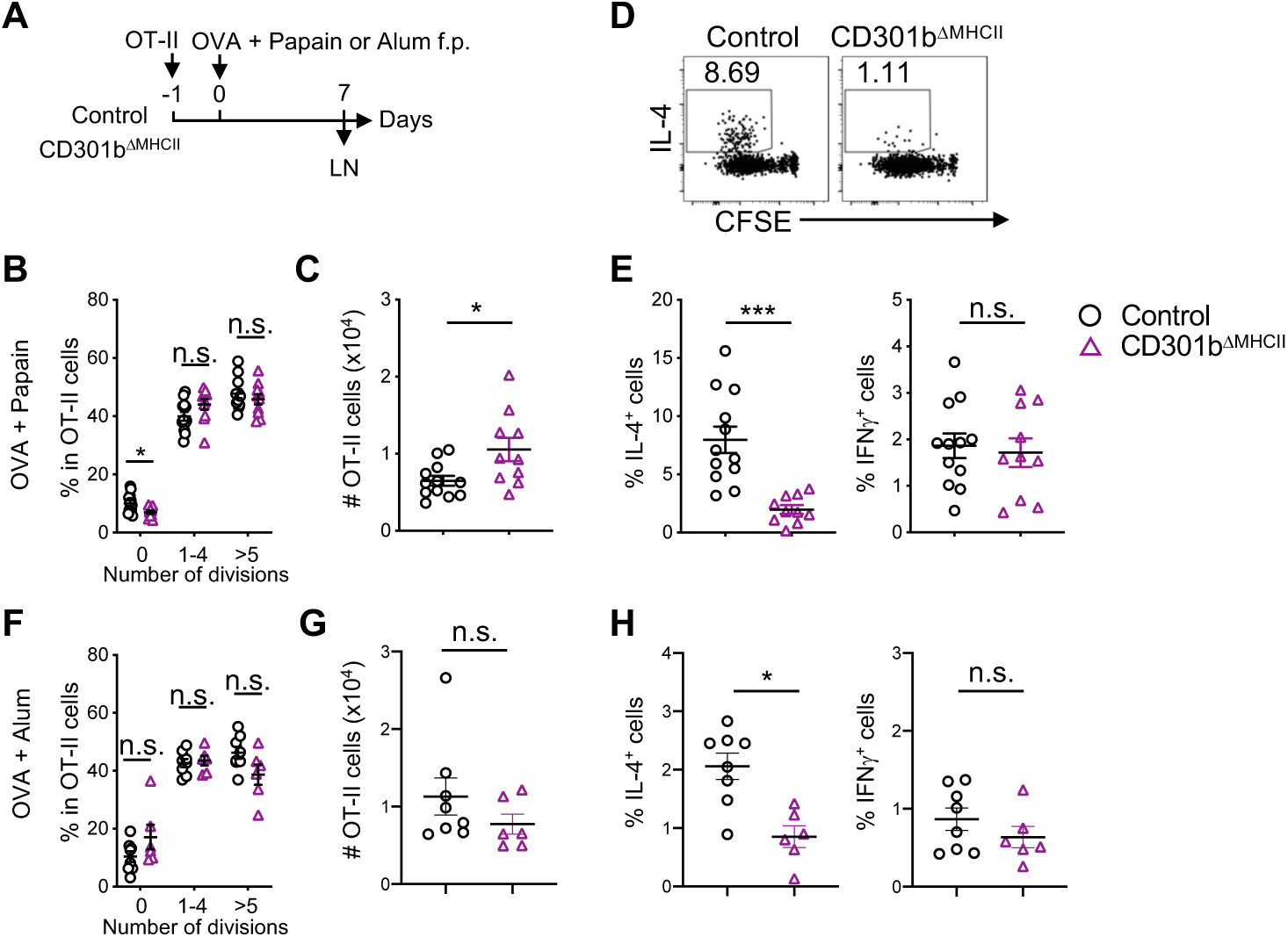
Direct antigen presentation by CD301b^+^ DCs is required for Th2 cell differentiation. **A**. Experimental design. Control (*Mgl2^+/Cre^*) and CD301b^ΔMHCII^ mice were adoptively transferred with 1×10^5^ CFSE-labeled OT-II cells and immunized one day later with OVA plus papain **(B-E)** or OVA plus alum **(F-H)** in the footpad. The draining popliteal LNs were harvested 7 days after the immunization and restimulated *ex vivo* with PMA and ionomycin. **B-H.** Frequencies of OT-II cells that have undergone indicated number of cell divisions **(B, F)**, numbers of OT-II cells **(C, G)**, flow cytometry plots of IL-4 and CFSE **(D)**, and frequencies of IL-4^+^ and IFNγ^+^ cells among the donor OT-II cells **(E, H)** are shown. Data represent means ± SEM of 12 control and 10 CD301b^ΔMHCII^ mice for **(B-E)**, 8 control and 6 CD301b^ΔMHCII^ mice for **(F-H)** or show representative flow cytometric plot of at least two independent experiments **(D)**. Not significant (n.s.), *p*≥0.05; \**p*<0.05; and \*\*\**p*< 0.001 by two-tailed Student’s t test.

### CD301b^+^ DCs are required for inducing full CD25 expression in CD4 T cells

DCs send critical activation signals to antigen-specific naive CD4T cells, leading to cell cycle entry and rapid upregulation of early activation markers such as CD69 and CD25 in T cells. We have shown previously that the depletion of CD301b^+^ DCs results in an 8-to 16-hour delay in CD69 upregulation and cell cycle entry of antigen-specific CD4T cells ^18^, but the link between the delayed activation and the selective impairment in Th2 differentiation remains unclear. To gain further insight on the potential defect in CD4T cell activation in these mice, we transferred CFSE-labeled OT-II cells into DT-treated WT (CD301b^+^ DC-intact) or Mgl2-DTR (CD301b^+^ DC-depleted) mice 24 hours after immunization with OVA plus papain in the footpad, and then blocked further LN entry of T cells with anti-CD62L monoclonal antibody (mAb) 2 hours later (**Fig. 2A**), which allows us to monitor the activation kinetics of OT-II cells in a synchronized manner ^18^. In agreement with our previous report ^18^, the upregulation of CD69 in OT-II cells was delayed in CD301b^+^ DC-depleted mice but was quickly recovered presumably due to antigen presentation by CD301b^−^ DCs (**Fig. 2B**). In contrast,

**Fig. 2.**
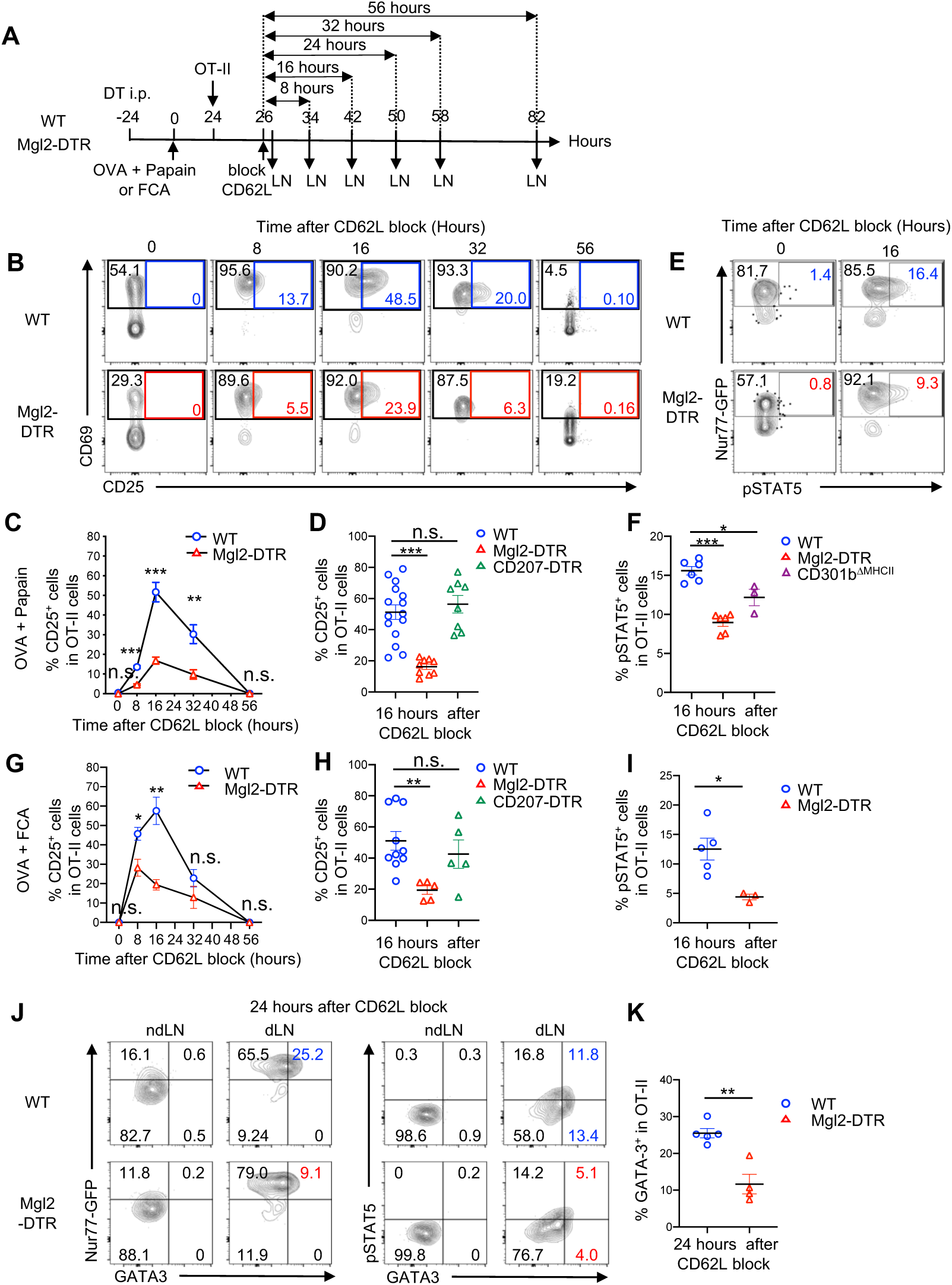
CD301b^+^ DCs are required for maximal CD25 expression and STAT5 activation in antigen-specific CD4T cells. **A**. Experimental design. Two million OT-II cells were transferred into DT-treated Mgl2-DTR or WT mice 24 hours after immunization with OVA plus papain **(B-F, J, K)** or OVA plus FCA **(G-I)** in the footpad. Two hours later, further homing of lymphocytes to LNs was blocked by injecting anti-CD62L mAb retro-orbitally. The dLNs were harvested at indicated time-points after the CD62L blockade. The donor OT-II cells were labeled by a congenic marker (CD45.1) or a cell tracer dye. OT-II cells expressing the Nur77-GFP reporter were used in **(E**, **F**, **I-K)**. DT-treated CD207-DTR mice and CD301b^ΔMHCII^ mice (without DT) were used as additional groups of recipients where indicated. **B-K.** Representative flow cytometric plots for indicated markers **(B, E, J)**, frequencies of CD25^+^ cells **(C, D, G, H)**, pSTAT5^+^ cells **(F, I)**, and GATA-3^+^ cells **(K)** among the donor OT-II cells are shown. Data represent means ± SEM **(C, D, F-I, K)** or show representative flow cytometry plots of at least two independent experiments **(B, E, J)**. In **(C)** and **(G)**, data are pooled from 4 to 12 mice per group at each time point. Not significant (n.s.), *p*≥0.05; \**p*<0.05; \*\**p*<0.01; and \*\*\**p*<0.001, by two-tailed Student’s t test.

CD25 expression was induced in a subset of CD69^+^ OT-II cells at a similar timing and peaked at 16 hours after the dLN entry in both CD301b^+^ DC-intact and CD301b^+^ DC-depleted mice, but with significantly lower expression levels in the latter (**Fig. 2B,C**). The requirement of CD301b^+^ DCs for the full CD25 upregulation is subset-specific, as the depletion of CD207^+^ DCs, including LCs and dermal CD103^+^ cDC1s ^20, 21, 22^, in CD207-DTR mice ^23^ did not affect CD25 expression (**Fig. 2D**).

CD25 (IL-2Rα) is the high affinity subunit of the IL-2R complex. Intracellular staining of phosphorylated STAT5 (pSTAT5) in OT-II cells expressing the Nur77-GFP reporter (Nur77-GFP;OT-II), whose expression correlates with the TCR signal strength ^24^, showed a reduction of pSTAT5, the major signaling component of IL-2R, in OT-II cells primed in CD301b^+^ DC-depleted mice. (**Fig. 2E, F**). In contrast, like CD69, expression of the Nur77-GFP reporter soon after the dLN entry was lower in Mgl2-DTR than in WT mice (**Fig. 2E**, 0h), but the comparable expression at a later time-point indicated similar amount of TCR stimulation received by OT-II cells over time regardless of the CD301b^+^ DC depletion status (**Fig. 2E**, 16h). The pSTAT5 levels were also reduced in CD301b^ΔMHCII^ mice (**Fig. 2F**), indicating the requirement of cognate interaction with CD301b^+^ DC for optimal activation of STAT5 in OT-II cells. Furthermore, the subset-specific requirement of CD301b^+^ DCs for the full CD25 upregulation and STAT5 phosphorylation in OT-II cells was also observed when the mice were immunized under a non-Th2 condition with Freund’s complete adjuvant (FCA) (**Fig. 2G-I**). The expression of GATA-3, the master regulator of Th2 cell differentiation, was induced in a subset of Nur77-GFP^+^ OT-II cells and showed positive correlation with pSTAT5 levels in the dLN of WT mice (**Fig. 2J**). The depletion of CD301b^+^ DCs resulted in a reduction of GATA-3 in OT-II cells in association with the impaired activation of STAT5 (**Fig. 2J,K**). Collectively, these data indicate that CD301b^+^ DCs deliver a qualitatively distinct signal required for the optimal CD25 upregulation in antigen-specific CD4T cells.

### Th2 cell differentiation relies more stringently on IL-2R signaling than Th1 cells

IL-2 plays fundamental roles in CD4T cell biology ^25, 26^. While some studies have suggested requirement of IL-2R signaling for Th2 cell differentiation *in vitro* ^27, 28^, its role in Th2-specific fate decision *in vivo* remains elusive, as abrogating IL-2R signaling often results in a broader defect in proliferation, differentiation and/or survival of CD4T cells ^29, 30^. To definitively address the role of IL-2R signaling in Th2 cell differentiation *in vivo*, we first neutralized IL-2 by intraperitoneally (i.p.) injecting an anti-IL-2 mAb S4B6-1, which blocks the binding of IL-2 to CD25 ^31, 32^, in WT mice immunized with OVA plus papain as in **Fig. 3A**. While the IL-2 blockade had no major impact on cell divisions and induced only a modest decrease in numbers of OT-II cells (**Fig. 3B,C**), it inhibited the differentiation of IL-4-producing Th2 cells without affecting their ability to induce IFNγ-producing Th1 cells (**Fig. 3D**), suggesting that IL-2 is specifically required for Th2 differentiation. Notably, the selective requirement of IL-2 for Th2 differentiation is not specific to the type 2 immune environment, since, when treated with the anti-IL-2 mAb, mice transferred with OT-II cells expressing the IL-4-GFP reporter (4get;OT-II) and immunized with OVA plus FCA, which predominantly induces non-type 2 inflammation, also exhibited significantly reduced IL-4-GFP expression in the donor 4get;OT-II cells without major alteration in their proliferation or IFNγ production (Supplementary **Fig. S1A-C**).

**Fig. 3.**
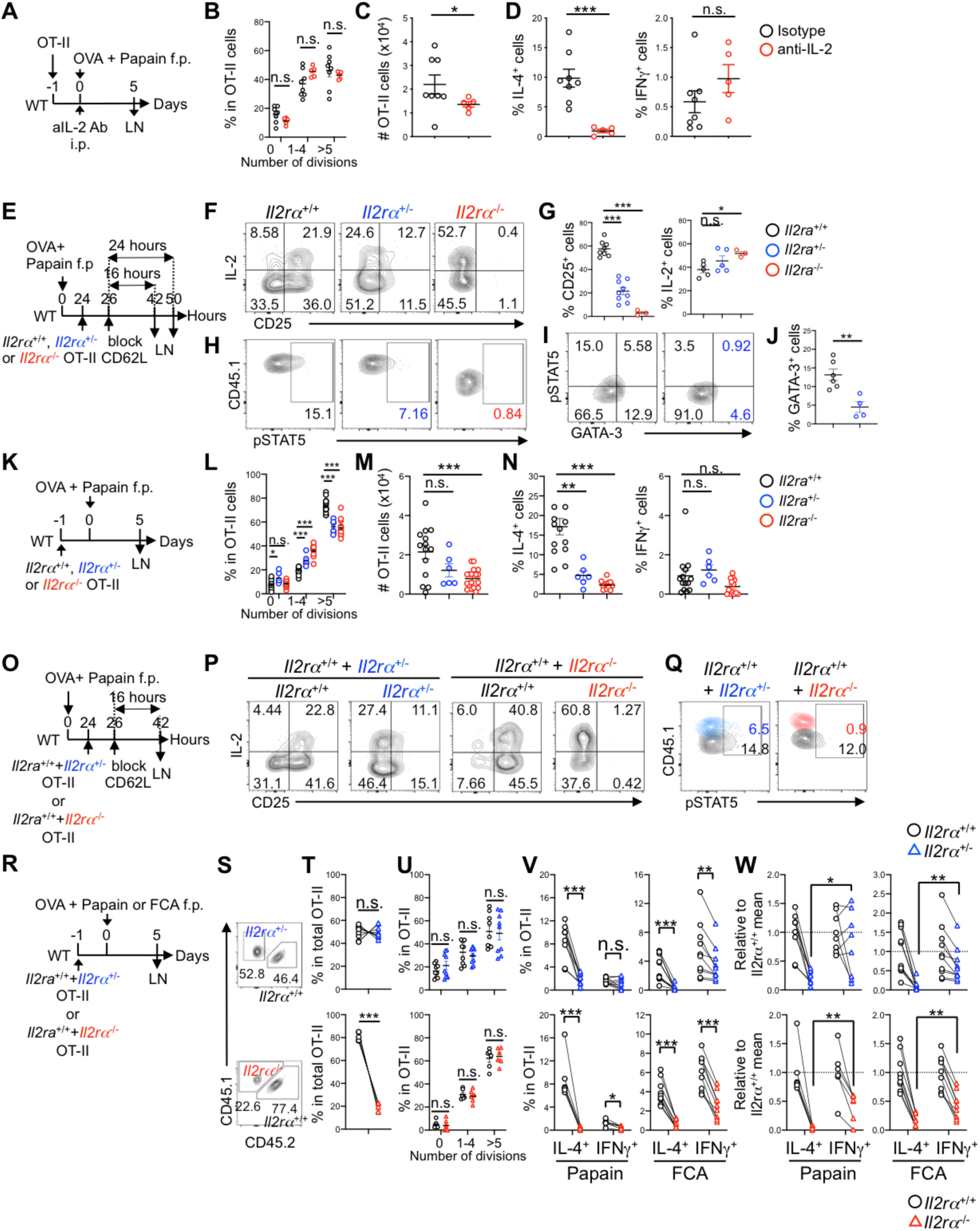
Th2 cell differentiation relies more stringently on IL-2R signaling than Th1 cells. **A-D**. WT mice were transferred with 1×10^5^ CFSE-labeled OT-II cells and immunized one day later with OVA plus papain in the footpad and injected i.p. with anti-IL-2 neutralizing mAb S4B6-1. The dLNs were harvested 5 days after the immunization and restimulated *ex vivo* with PMA and ionomycin **(A)**. Frequencies of OT-II cells that have undergone indicated number of cell divisions **(B)**, numbers of OT-II cells **(C)**, and frequencies of IL-4^+^ and IFNγ^+^ cells among the donor OT-II cells **(D)** are shown. **E-J.** CFSE-labeled *Il2ra^+/+^*, *Il2ra^+/−^*, or *Il2ra^−/−^* OT-II (2×10^6^) cells were transferred into WT mice 24 hours after immunization with OVA plus papain in the footpad and allowed to home to LNs for 2 hours, after which further homing was blocked with anti-CD62L mAb **(E)**. The dLNs were harvested 16 **(F-H)** or 24 hours **(I, J)** after the CD62L blockade. For intracellularly staining IL-2, cells were restimulated *ex vivo* with PMA and ionomycin. Representative flow cytometric plots of cell surface CD25 and intracellular IL-2 **(F)**, pSTAT5 **(H, I)**, or GATA-3 **(I)** on the donor OT-II, frequencies of CD25^+^ and IL-2^+^ cells **(G)** and GATA-3^+^ cells **(J)** among the donor OT-II cells are shown. **K-N.** WT mice were transferred with either *Il2ra^+/+^*, *Il2ra^+/−^*, or *Il2ra^−/−^*CFSE-labeled 1×10^5^ OT-II cells separately, and immunized one day later with OVA plus papain in the footpad. The dLNs were harvested 5 days after the immunization and restimulated in vitro with PMA and ionomycin **(K)**. Frequencies of OT-II cells that have undergone indicated number of cell divisions **(L)**, numbers of OT-II cells **(M)**, and frequencies of IL-4^+^ and IFNγ^+^ cells among the donor OT-II cells **(N)** are shown. **O-Q.** As in **(E)**, but a 1:1 mixture (1×10^5^ cells each) of *Il2ra^+/+^* (CD45.1/.2) and *Il2ra^+/−^*(CD45.1/.1) OT-II cells, or *Il2ra^+/+^* (CD45.1/.2) and *Il2ra^−/−^* OT-II (CD45.1/.1) cells were co-transferred. The dLNs were harvested 16 hours after the CD62L blockade. Representative flow cytometry plots of CD25 and IL-2 **(P)** and pSTAT5 **(Q)** among the donor OT-II cells are shown. **R-W.** WT mice were co-transferred with a 1:1 mixture (1×10^5^ cells each) of *Il2ra^+/+^* (CD45.1/.2) and *Il2ra^+/−^* (CD45.1/.1) OT-II cells, or *Il2ra^+/+^*(CD45.1/.2) and *Il2ra^−/−^* (CD45.1/.1) OT-II cells, and immunized one day later with OVA plus papain in the footpad. The dLNs were harvested 5 days after the immunization and restimulated *ex vivo* with PMA and ionomycin **(R)**. Flow cytometry plots of CD45.1 and CD45.2 expression by gated CD45.1⁺ OT-II cells **(S)**, paired frequencies of indicated OT-II cells among the total OT-II cells within individual host dLNs **(T)**, frequencies of indicated OT-II cells that have undergone indicated number of cell divisions **(U)**, paired relative frequencies of IL-4^+^ and IFNγ^+^ cells among the CD44^+^ CFSE^lo^ OT-II cells within individual host dLNs in response to OVA and papain (Left) or FCA (Right) **(V)**. In **(W)**, frequencies of IL-4^+^ and IFNγ^+^ of *Il2ra^+/−^* or *Il2ra^−/−^* OT-II cells in **(V)** were normalized to the mean frequency of the *Il2ra^+/+^* OT-II counterpart. In **(T, V, W)**, each connecting line indicates OT-II cells of two different genotypes isolated from the same host. Data represent means ± SEM **(B-D, G, J, L-N, T-W)** or show representative flow cytometry plots of at least two independent experiments **(F, H, I, P, Q, S)**. Not significant (n.s.), *p*≥0.05; \**p*<0.05; \*\**p*< 0.01; and \*\*\**p*<0.001, by two-tailed Student’s t test **(B-D, G, J, L-N, U)** or paired t test **(T, V, W)**.

Since the anti-IL-2 mAb S4B6-1 can either neutralize or potentiate IL-2R signaling in a dose-dependent manner ^33, 34^, we next examined if the reduced, but not completely abrogated, CD25 upregulation in CD4T cells was responsible for the impaired Th2 differentiation in CD301b^+^ DC-depleted mice by transferring *Il2ra* (CD25)^+/+^, *Il2ra*^+/−^, or *Il2ra*^−/−^ OT-II cells into WT mice immunized with OVA plus papain (**Fig. 3E**). As expected, CD25 expression in the *Il2ra*^+/−^ OT-II cells was reduced by approximately 50% when measured 16 hours after the dLN entry, whereas IL-2 production was increased inversely with the *Il2ra* gene dosage likely due to the lack of the CD25-dependent negative feedback ^35^ (**Fig. 3F,G**). Nevertheless, STAT5 phosphorylation was reduced by roughly 50% and nearly absent in the *Il2ra*^+/−^ and *Il2ra*^−/−^ OT-II cells, respectively (**Fig. 3H**), suggesting that the CD25 expression level, rather than the IL-2 production, from OT-II cells is the primarily determinant of STAT5 phosphorylation in this model. The reduction of pSTAT5 in the *Il2ra*^+/−^

OT-II cells was associated with lower levels of GATA-3 expression 24 hours after the priming (**Fig. 3I,J**), indicating that the maximal IL-2R signaling is necessary for the early commitment to the Th2 cell fate. Five days after immunization with OVA plus papain without CD62L blockade (**Fig. 3K**), >80% of *Il2ra*^+/−^ and *Il2ra*^−/−^ OT-II cells underwent cell divisions, though there were mild defects in the number of division cycles as well as the total numbers of the donor OT-II cells (**Fig. 3L,M**). However, the divided *Il2ra*^+/−^ and *Il2ra*^−/−^ OT-II cells failed to produce IL-4 without showing a defect in IFNγ production (**Fig. 3N**). Immunization with OVA plus FCA also resulted in a loss of IL-4 production in the *Il2ra*^+/−^ OT-II cells, but in this case with a trend for reduction in IFNγ production without significant impact on proliferation (Supplementary **Fig. S1D-F**).

While the above data suggest Th cell-intrinsic requirement of full CD25 expression for Th2 cell differentiation, the potentially elevated IL-2 levels in the dLN due to the increased production from the *Il2ra*^+/−^ and *Il2ra*^−/−^ OT-II cells (**Fig. 3F,G**) may have activated bystander T regulatory (Treg) cells and indirectly suppressed Th cell differentiation as previously shown for self-reactive Th cells ^36, 37^. To further elucidate the role of IL-2R signaling in Th2 fate decision *in vivo*, we next co-transferred equal numbers of *Il2ra*^+/+^ and *Il2ra*^+/−^ or *Il2ra*^−/−^ OT-II cells so that the IL-2 availability in the dLN is normalized (**Fig. 3O,R**). The *Il2ra* gene dosage-dependent decrease and increase in IL-2 production and STAT5 phosphorylation, respectively, was also observed in this setting (**Fig. 3P,Q**), but, unlike the individual transfer approach (**Fig. 3L**), the *Il2ra*^+/−^ OT-II cells showed comparable expansion to the co-transferred *Il2ra*^+/+^ counterpart regardless of the adjuvant used, whereas the *Il2ra*^−/−^ OT-II cells were outcompeted significantly by the *Il2ra*^+/+^ OT-II cells (**Fig. 3S-U, Supplementary Fig. S1G-I**). Importantly, the loss of even a single copy of *Il2ra* in the donor OT-II cells consistently resulted in a dramatic reduction in IL-4 production in a manner independent of the degree of OT-II cell expansion, whereas its impact on IFNγ production was variable and less significant (**Fig. 3V,W**). Collectively, these data indicate that the Th2 fate decision by antigen-specific CD4 T cells *in vivo* requires a potent and cell-intrinsic IL-2R signaling and is more sensitive to a partial loss of CD25 than the Th1 cell differentiation.

### CD301b^+^ DC-derived IL-2 is required for the full CD25 upregulation and Th2 differentiation in antigen-specific CD4T cells

Following TCR ligation, CD25 is induced in antigen-specific CD4T cells through two consecutive steps: the initial upregulation induced by the TCR signaling and the subsequent TCR-independent amplification of the *Il2ra* transcription by the IL-2-driven, CD25-dependent positive feedback ^38, 39^. While the activated CD4T cells themselves are generally considered as the critical source of IL-2 in this feedback loop ^39, 40, 41^, DCs can also produce IL-2 upon activation in both mice and humans ^42, 43, 44, 45, 46^, though its role in T cell priming *in vivo* remains elusive. In mice immunized with OVA plus papain (**Fig. 4A**), we found that CD301b^+^ DCs expressed higher levels of IL-2 than CD301b^−^ DCs or than those in the non-draining lymph node (ndLN) (**Fig. 4B,C**). To examine if CD301b^+^ DC-derived IL-2 plays a critical role in CD25 upregulation and Th2 fate decision in antigen-specific CD4T cells, we generated mice lacking IL-2 specifically in CD301b^+^ DCs (*Mgl2^+/cre^*;*Il2^fl/fl^*, CD301b^ΔIL-2^ mice) ^47^ (Supplementary **Fig. S2A,B**) and monitored the activation and differentiation of the donor OT-II cells upon immunization with OVA plus papain (**Fig. 4D-L**). Notably, blocking IL-2 in WT recipients resulted in a reduced CD25 expression in the donor OT-II cells. Likewise, the donor OT-II cells failed to fully upregulate CD25 in CD301b^ΔIL-2^ mice even though nearly all the OT-II cells expressed CD69 (**Fig. 4E-G**), indicating the specific requirement of CD301b^+^ DC-derived IL-2 for fully upregulating CD25 in antigen-specific CD4T cells. Furthermore, the differentiation of Th2 cells, monitored using the 4get;OT-II reporter cells, was abolished in the CD301b^ΔIL-2^ recipients without affecting the cell cycle progression, expansion, or their differentiation into IFNγ-producing Th1 cells (**Fig. 4H-L**). Taken together, these results demonstrate that CD301b^+^ DC-derived IL-2 is critical for inducing maximal CD25 upregulation and Th2 fate decision by antigen-specific CD4T cells.

**Fig. 4.**
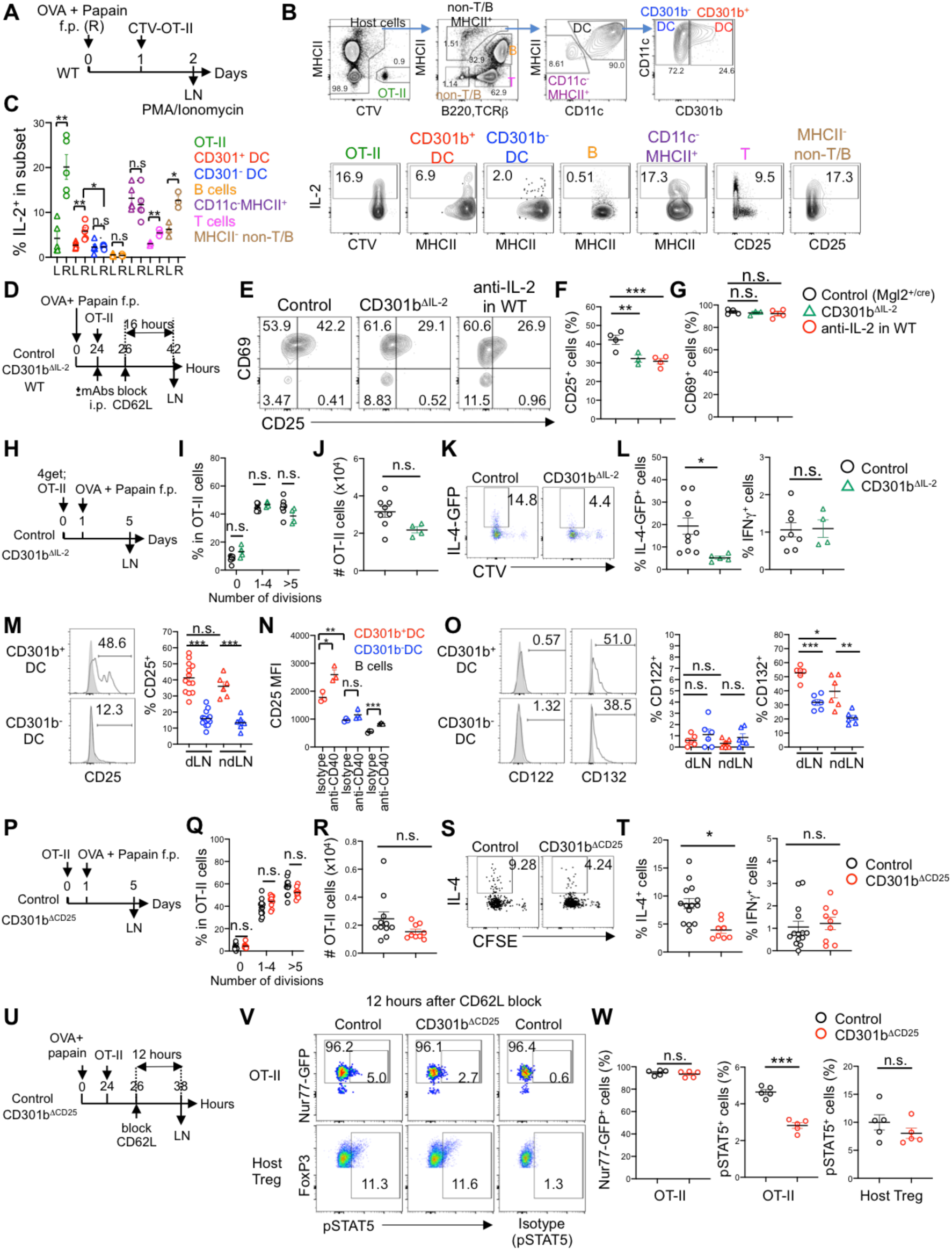
CD301b^+^ DC-derived IL-2 and CD25 are required for the maximal IL-2R signaling and Th2 cell differentiation of antigen-specific CD4T cells. **A-C**. WT mice were immunized with OVA plus papain in the right (R) footpad and adoptively transferred one day later with 1×10^6^ Cell Tracer Violet (CTV)-labeled OT-II cells. The dLNs (R) and left (L) ndLNs were harvested 24 hours after the transfer and restimulated *ex vivo* with PMA and ionomycin for intracellularly staining IL-2 **(A)**. Frequencies of IL-2-producing cells in each cell type were defined as in **(B)** and enumerated **(C)**. **D-G.** Two million CFSE-labeled OT-II cells were transferred into control (*Mgl2^+/Cre^*), CD301b^ΔIL-2^ or WT mice 24 hours after immunization with OVA plus papain in the footpad and allowed to home to LNs for 2 hours, after which further homing was blocked with anti-CD62L mAb. The WT recipients were also treated with anti-IL-2 neutralizing mAb. The dLNs were harvested 16 hours after the CD62L blockade **(D)**. Representative flow cytometry plots of CD69 and CD25 **(E)**, frequencies of CD25^+^ **(F)** and CD69^+^ **(G)** cells among the donor OT-II cells in the dLN are shown. **H-L.** Control and CD301b^ΔIL-2^ mice were transferred with 1×10^5^ CTV-labeled 4get;OT-II cells and immunized one day later with OVA plus papain in the footpad. The dLNs were harvested 5 days after the immunization and restimulated *ex vivo* with PMA and ionomycin. Frequencies of OT-II cells that have undergone indicated number of cell divisions **(I)**, numbers of OT-II cells **(J)**, representative flow cytometry plots of IL-4-GFP reporter and CTV dilution **(K)**, and frequencies of IL-4-GFP^+^ Th2 and IFNγ^+^ Th1 cells among the donor OT-II cells **(L)** are shown. **M-O**. Expression of IL-2R subunits in CD301b^+^ and CD301b^−^ DCs. WT mice were immunized with OVA plus papain in the footpad **(M,O)**, or i.p. injected with anti-CD40 agonistic mAb or an isotype control mAb **(N)**. The popliteal LNs were harvested 24 hours after the immunization or mAb injection for analyzing the expression of CD25 **(M,N)**, CD122 and CD132 **(O)** in indicated cell types. Gray histograms indicate the binding of respective isotype control mAb. **P-T.** Control (*Mgl2^+/Cre^*) and CD301b^ΔCD^^25^ mice were transferred with 1×10^5^ CFSE-labeled OT-II cells and immunized one day later with OVA plus papain in the footpad **(P)**. The dLNs were harvested 5 days after the immunization and restimulated *ex vivo* with PMA and ionomycin. Frequencies of OT-II cells that have undergone indicated number of cell divisions **(Q)**, numbers of OT-II cells **(R)**, representative flow cytometry plots of IL-4 and CFSE dilution **(S)**, and frequencies of IL-4^+^ Th2 and IFNγ^+^ Th1 cells among the donor OT-II cells **(T)** are shown. **U-W.** Two million CTV-labeled Nur77-GFP;OT-II cells were transferred into control or CD301b^ΔCD^^25^ mice 24 hours after immunization with OVA plus papain in the footpad and allowed to home to LNs for 2 hours, after which further homing was blocked with anti-CD62L mAb. The dLNs were harvested 12 hours after the CD62L blockade **(U)**. Representative flow cytometry plots of pSTAT5 and Nur77-GFP in OT-II cells or pSTAT5 and FoxP3 among CD25^+^ FoxP3^+^ CD4^+^ Treg cells **(V)**, frequencies of Nur77-GFP^+^ or pSTAT5^+^ among OT-II cells, and pSTAT5^+^ among Treg cells **(W)** in the dLN are shown. Data represent means ± SEM **(C, F, G, I, J, L, M-O, Q, R, T, W)**, or show representative flow cytometry plots of at least two independent experiments **(B, E, K, M, O, S, V)**. Not significant (n.s.), *p*≥0.05; \**p*<0.05; \*\**p*<0.01; and \*\*\**p*<0.001, by two-tailed Student’s t test **(C, F, G, I, J, L, M-O, Q, R, T, W)**.

### CD301b^+^ DC-derived CD25 is required for Th2 cell differentiation

The above data suggest that CD301b^+^ DCs provide IL-2 specifically to cognate CD4 T cells, but it remains unclear how the directed action of IL-2 is protected from diffusion, which would lead to rapid consumption of IL-2 by bystander Treg cells rather than by cognate CD4T cells ^36, 37, 40, 41, 48^. Previous studies have suggested that the diffusion of IL-2 is limited by the expression of CD25 on non-target cells ^45, 49, 50, 51^. We found that the expression of CD25 was higher in CD301b^+^ DCs than in CD301b^−^ DCs and was also enhanced upon CD40 stimulation (**Fig. 4M,N**). CD301b^+^ DCs also expressed higher levels of CD132 (IL-2Rγ) than CD301b^−^ DCs, but they both express little, if any, CD122 (IL-2Rβ) (**Fig. 4O**), suggesting that the IL-2R in CD301b^+^ DCs does not efficiently signal, as CD122 is required for the signaling ^26^. Importantly, in mice lacking CD25 specifically in CD301b^+^ DCs (*Mgl2^+/Cre^;Il2ra^fl/fl^*, CD301b^ΔCD25^ mice) ^52, 53^ (Supplementary **Fig. S2C,D**), differentiation of the donor OT-II cells into Th2 cells was significantly impaired without showing a defect in their expansion or Th1 cell differentiation (**Fig. 4P-T**). The lack of CD25 in CD301b^+^ DCs resulted in a reduction of pSTAT5 specifically in the donor OT-II cells but not in the endogenous Treg cells (**Fig. 4U-W**), suggesting that CD301b^+^ DC-intrinsic CD25 expression facilitates the directed action of IL-2 toward cognate CD4T cells. Collectively, these results indicate that CD301b^+^ DCs promote Th2 cell differentiation through reinforcing the availability of IL-2 for cognate CD4 T cells by the DC-intrinsic expression of CD25.

### CD40 stimulation induces IL-2 production specifically by CD301b^+^ DCs

To further explore the mechanism for the Th2 cell fate instruction by CD301b^+^ DCs, we performed CITEseq, a single-cell RNA sequencing (scRNAseq) technique with DNA-barcoded mAbs for simultaneously analyzing the expression of mRNA and cell surface markers ^54^, on MHCII^hi^ migratory DCs isolated form skin-dLNs of naïve and papain-immunized mice one day after immunization. Unsupervised clustering analysis of the cell surface markers identified 7 distinct clusters composed of 3 major DC subsets, including LC (cluster 4: CD172a^lo^ CD326^hi^ XCR1^−^), cDC1s (cluster 5: CD172a^−^ CD326^int^ XCR1^+^), and cDC2s (clusters 1, 2, 3, 6 and 7: CD172a^hi^ CD326^−^ XCR1^−^) (**Fig. 5A**). The cDC2s were divided into three subpopulations, CD301b^lo^, CD301b^int^, or CD301b^hi^ DCs, based on CD301b expression levels (**Fig. 5B**). Alternatively, the cDC2s were further divided into five subclusters based on their mRNA expression profile, with the subclusters 2 and 5 mainly composed of naive cDC2s, while subclusters 1, 3, and 4 predominantly derived from cDC2s in immunized mice (**Fig. 5A,C**). While the CD301b expression alone did not identify a single cDC2 subcluster in either naive or immunized mice, cells in the cDC2 subcluster 1, 2 and 3 were more enriched in CD301b^int^ and CD301b^hi^ DCs, whereas those in the subcluster 4 and 5 were more abundant in CD301b^lo^ DCs (**Fig. 5B,C**). The Ingenuity Pathway Analysis ^55^ of the genes differentially expressed in CD301b^hi^ DCs compared with all other DC subsets in papain-immunized mice identified several pathways including the CD40 pathway, which was enriched in CD301b^hi^ DCs in both naive and papain-immunized mice (**Fig. 5D,E**). Indeed, CD301b^+^ DCs expressed higher levels of CD40 compared to CD301b^−^ DCs in both dLN ndLN of papain-immunized mice (**Fig. 5F,G**). Importantly, stimulation of CD40 by i.p. administration of an agonistic anti-CD40 mAb FGK4.5 ^56, 57, 58^ induced IL-2 production in CD301b^+^ DCs along with CD80 and CD86 (**Fig. 5H,I**). Notably, the CD40-induced IL-2 production was specific to CD301b^+^ DCs among DC subsets (**Fig. 5H**). Thus, CD40 ligation stimulates IL-2 production and upregulation of costimulatory molecules in CD301b^+^ DCs.

**Fig. 5.**
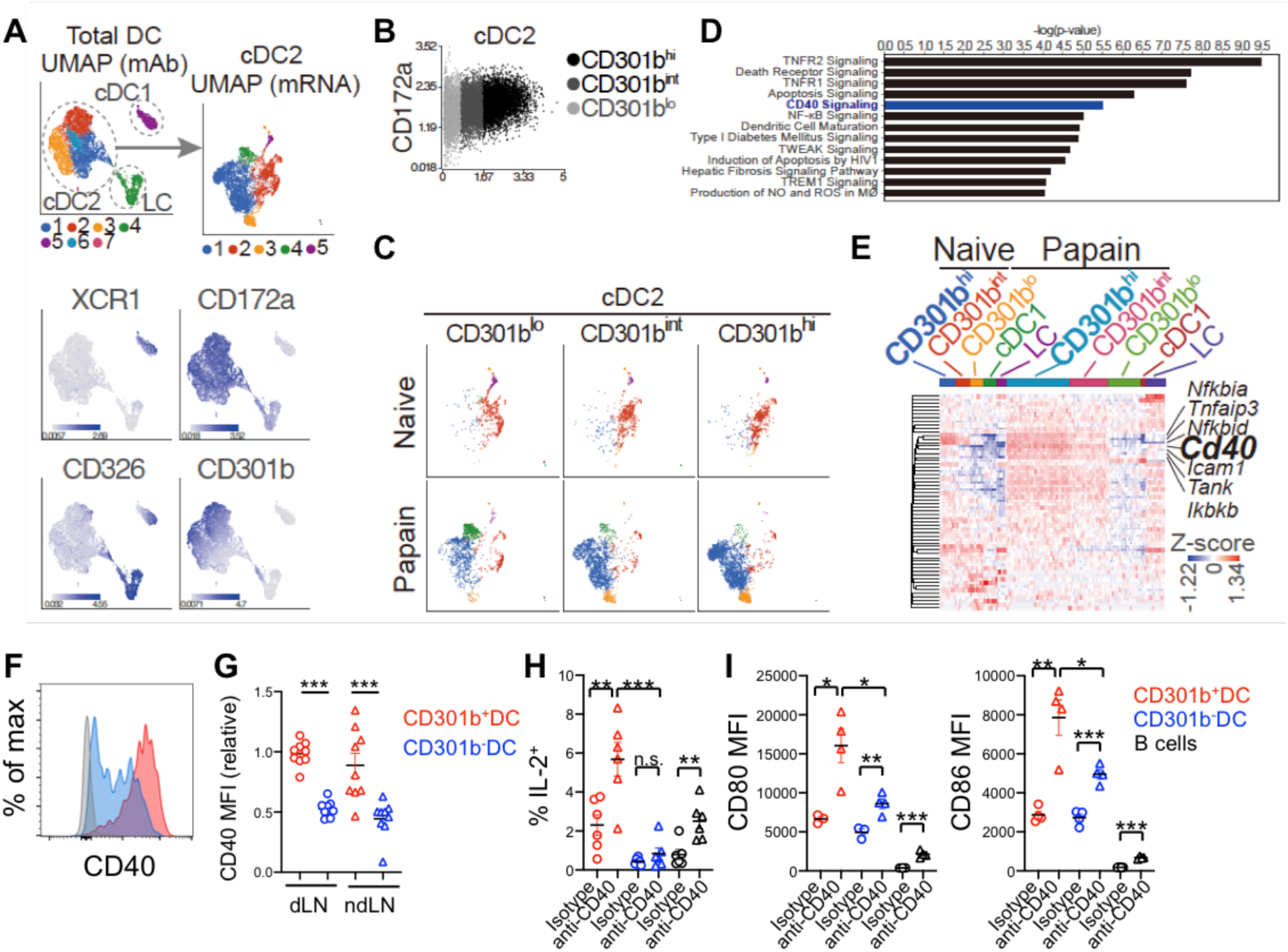
CD40 stimulation induces IL-2 production by CD301b+ DCs. **A-E**. MHCII^hi^ migratory DCs (including LCs) were sorted from the skin-dLNs of naïve mice and mice immunized with OVA plus papain for one day and labeled with DNA-barcoded mAbs for CD8α, CD11b, CD11c, CD24, CD64, CD103, CD172a, CD301b, CD326, Ly6C, MHCII and XCR1 for CITEseq analysis. Uniform Manifold Approximation and Projection (UMAP) plots for the seven distinct DC clusters separated by the cell surface marker expression (left) and the five cDC2 subclusters identified based on the mRNA expression (right) are shown **(A)**. Clusters of cDC1, cDC2 and LCs were identified by the expression of XCR1, CD172a and CD326, respectively, as indicated. cDC2 cells were divided into 3 subpopulations based on the cell surface CD301b expression as in **(B)** and projected onto the UMAP plots **(C)**. Ingenuity Pathway Analysis identified the CD40 pathway as one of the differentially expressed pathways in CD301b^hi^ cDC2s compare with all other DC populations in papain-immunized dLNs **(D)**. The heatmap in **(E)** shows the expression of each gene in the CD40 pathway in each sequenced cell. **F, G.** Expression of CD40 in CD301b^+^ and CD301b^−^ DCs in the dLN and ndLN of WT mice immunized with papain in the footpad for one day. Flow cytometry histograms in the dLN **(F)** and mean fluorescence intensity (MFI) of CD40 **(G)** are shown. Gray histogram in F shows the binding of an isotype control mAb. **H, I.** WT mice were i.p. injected with anti-CD40 agonistic or isotype control mAbs. Popliteal LNs were harvested 24 hours after the injection and stimulated *ex vivo* with PMA and ionomycin for intracellularly staining IL-2. Alternatively, the cell surface expression of CD80 and CD86 was examined by flow cytometry. Frequencies of IL-2^+^ cells among indicated subsets **(H)** and MFI of CD80 and CD86 in indicated populations **(I)** are shown. Data represent means ± SEM **(G-I)**, or show representative flow cytometry plots of at least two independent experiments **(F)**. Not significant (n.s.), *p*≥0.05; \**p*<0.05; \*\**p*<0.01; and \*\*\**p*<0.001, by two-tailed Student’s t test **(G-I)**.

### CD301b^+^ DC-intrinsic CD40 is required for Th2 cell differentiation

Previous studies have shown the requirement of CD40 in Th2 cell differentiation ^59, 60^, but the underlying mechanism is unknown. Indeed, the 4get;OT-II cells primed with OVA plus papain in CD40-deficient mice expressed significantly lower levels of GFP (reporting *Il4*) ^61^ compared to those in WT mice, whereas their expansion, cell division, and differentiation into Th1 cells remained comparable (Supplementary **Fig. S3A-C**). Moreover, a similar reduction was observed for Th2 cells but not for Th1 cells when the OT-II cells were primed in mixed bone marrow (BM)-chimeric (BMC) mice reconstituted with a 1:1 mixture of MHCII-deficient and CD40-deficient BM cells, in which cells expressing MHCII do not express CD40, while cells expressing CD40 cannot present antigens to CD4T cells (**Fig. 6A-C**). Intriguingly, unlike OT-II cells primed in CD40-deficient mice, the cell division and expansion of OT-II cells was also impaired in MHCII^−/−^:CD40^−/−^ BMC mice (Supplementary **Fig. S3D,E**), suggesting that the partial loss of MHCII-dependent cognate TCR stimulation in the BMC mice had an additional negative impact on priming. These data indicate the requirement of the concurrent CD40 signaling and antigen presentation in the same cells specifically for Th2 fate instruction.

**Fig. 6.**
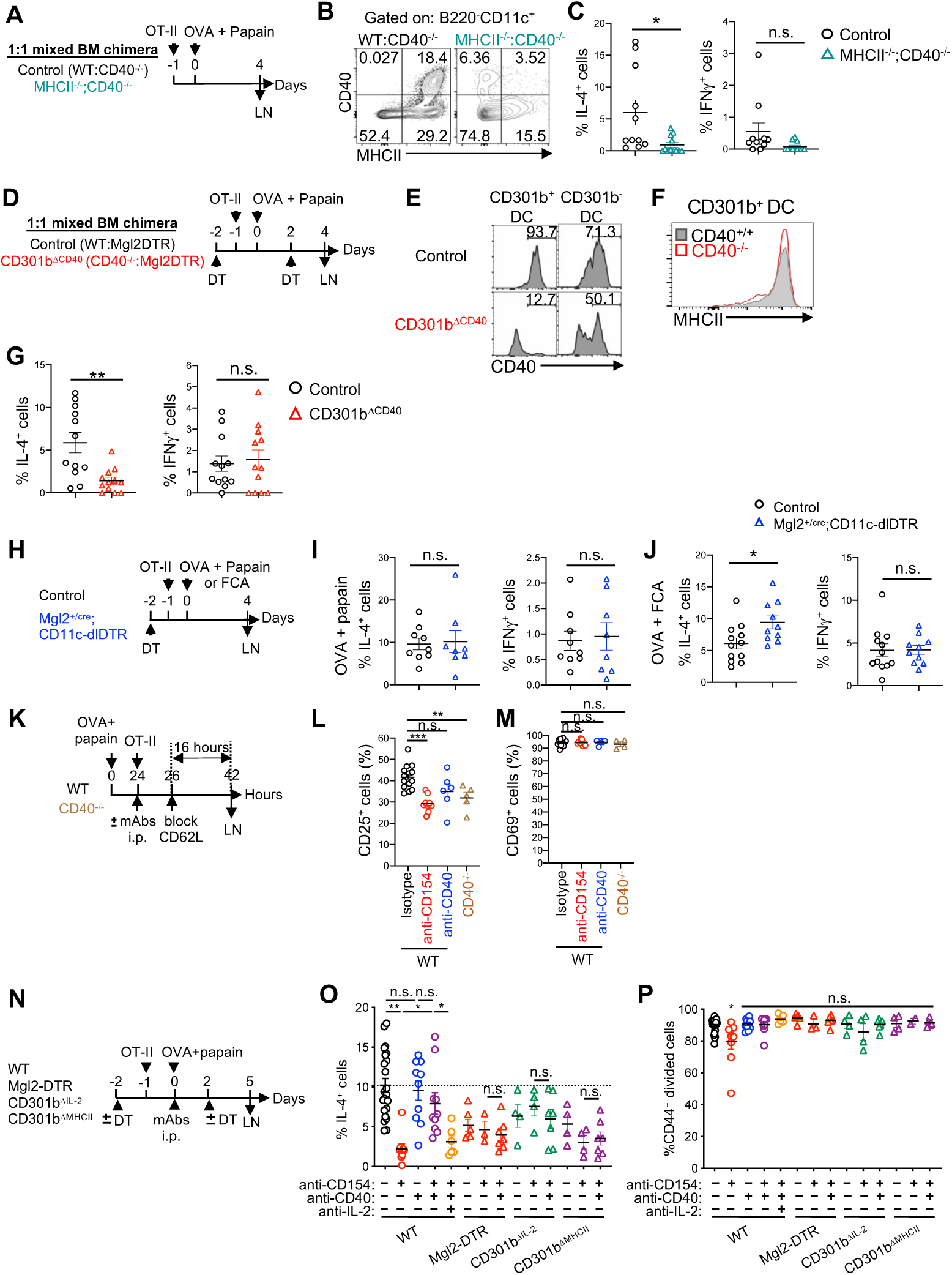
CD301b^+^ DCs instruct Th2 cell fate on CD4T cells through the DC-intrinsic CD40–IL-2 axis. **A-C**. Control (WT;CD40^−/−^) and MHCII^−/−^;CD40^−/−^ mixed BMC mice were transferred with 1×10^5^ CFSE-labeled OT-II cells and immunized one day later with OVA plus papain in the footpad. The dLNs were harvested 4 days after the immunization and restimulated *ex vivo* with PMA and ionomycin **(A)**. Representative flow cytometry plots of the expression of MHCII and CD40 in B220^−^ CD11c^+^ cells **(B)** and frequencies of IL-4^+^ and IFNγ^+^ cells among the donor OT-II cells in the dLN **(C)** are shown. **D-G.** Control and CD301b^ΔCD^^40^ mixed BMC mice were transferred with 1×10^5^ CFSE-labeled OT-II cells and immunized one day later with OVA plus papain in the footpad. The mice were treated with DT on days −2 and +2, and the dLNs were harvested 4 days after the immunization and restimulated *ex vivo* with PMA and ionomycin. Representative flow cytometry histograms of CD40 in CD301b^+^ and CD301b^−^ DCs of control and CD301b^ΔCD^^40^ mice **(E)**, MHCII expression in CD301b^+^ DCs of LNs from WT and CD40^−/−^ mice **(F)**, frequencies of IL-4^+^ and IFNγ^+^ cells among the donor OT-II cells in the dLN **(G)** are shown. **H-J.** DT-treated control (*Mgl2^+/cre^*) and *Mgl2^+/cre^*;CD11c-dlDTR mice were transferred with 1×10^5^ CFSE-labeled OT-II cells and immunized one day later with OVA plus papain or OVA plus FCA in the footpad on Day 0. The dLNs were harvested on day 4 and restimulated *ex vivo* with PMA and ionomycin **(H)**. Frequencies of IL-4^+^ and IFNγ^+^ cells among the donor OT-II cells in the dLN of mice immunized with OVA plus papain **(I)** and with OVA plus FCA **(J)** are shown. **K-M.** Two million CSFE-labeled OT-II cells were transferred into WT or CD40^−/−^ mice 24 hours after immunization with OVA plus papain in the footpad. WT mice were i.p. injected with indicated mAbs at the time of OT-II transfer. Two hours later, further homing of lymphocytes to LNs was blocked by injecting anti-CD62L mAb. The dLNs were harvested 16 hours after the CD62L blockade **(K)**. Frequencies of CD25^+^ **(L)** and CD69^+^ **(M)** cells among the donor OT-II cells in the dLN are shown. **N-P.** WT, Mgl2-DTR, CD301b^ΔIL-2^, and CD301b^ΔMHCII^ mice were transferred with 1×10^5^ CFSE-labeled OT-II cells and immunized one day later with OVA plus papain in the footpad and simultaneously injected i.p. with indicated mAbs. Mgl2-DTR mice were treated with DT on days –2 and +2. The dLNs were harvested 5 days after the immunization and restimulated *ex vivo* with PMA and ionomycin **(N)**. Frequencies of IL-4^+^ cells **(O)** and CFSE^lo^ CD44^+^ cells **(P)** among the donor OT-II cells are shown. Data represent means ± SEM **(C, G, I, J, L, M, O, P)**, or show representative flow cytometry plots of at least two independent experiments **(B, E, F)**. Not significant (n.s.), *p*≥0.05; \**p*<0.05; \*\**p*<0.01; and \*\*\**p*<0.001, by two-tailed Student’s t test **(C, G, I, J, L, M, O, P)**.

We next reconstituted lethally irradiated WT mice with a 1:1 mixture of CD40^−/−^ and Mgl2-DTR BM cells (**Fig. 6D**). In the resulting mice (CD301b^ΔCD40^), only CD40^−/−^ cells remained intact in the CD301b^+^ compartment upon DT injection, while all CD301b^−^ compartments maintained the 1:1 chimerism (**Fig. 6E**). Although the MHCII expression in CD40^−/−^ CD301b^+^ DCs was comparable to that in WT CD301b^+^ DCs (**Fig. 6F**), the OT-II cells primed with OVA plus papain in the CD301b^ΔCD40^ mice had significantly reduced Th2 cells but not Th1 cells (**Fig. 6G**). Similarly to the MHCII^−/−^:CD40^−/−^ BMC mice (Supplementary **Fig. S3D,E**), there was a slight but significant reduction in the cell division and expansion of OT-II cells in the CD301b^ΔCD40^ mice (Supplementary **Fig. S3F,G**), suggesting that the loss of 50% of CD301b^+^ DCs in these mice partially attenuated the priming itself. Collectively, these data indicate that CD301b^+^ DC-intrinsic CD40 is required for Th2 cell fate instruction.

### CD301b^+^ DCs are a sufficient DC subset for Th2 cell fate instruction

The data thus far indicate that direct antigen presentation, CD40 expression and IL-2 production by CD301b^+^ DCs are all *required* for Th2 cell fate instruction, but involvement of multiple DC subsets for Th cell differentiation has also been suggested ^62, 63^. To address if CD301b^+^ DCs are *sufficient* and no other DC subsets are required for Th2 cell fate instruction, we took advantage of mice expressing a Cre-removable DTR cassette in CD11c^+^ cells (CD11c-dlDTR) crossed with the Mgl2-Cre strain (Mgl2-Cre;CD11c-dlDTR mice), in which CD301b^+^ DCs are protected from DT-induced depletion while CD301b^−^ DCs are susceptible to DT ^18^. Although the number of CD301b^+^ cells in these mice are underestimated due to the loss of CD301b protein expression from the *Mgl2^cre^* allele ^18^, the use of the Cre-inducible Rosa26-loxP-stop-loxP (LSL)-tdTomato (iTom) reporter concurrently with these mice (Mgl2-Cre;CD11c-dlDTR;R26^LSL-iTom^) confirmed subset-specific protection of Cre-expressing *Mgl2* (CD301b)^+^ DCs from DT-induced depletion in the skin-dLNs (Supplementary **Fig. S3H-M**). The majority of the protected DCs expressed PD-L2, a marker known to be correlated with CD301b expression in CD172a^+^ cDC2s ^10, 17^, whereas all other DCs, including PD-L2^−^ cDC2s, LCs, and XCR1^+^ cDC1s, were effectively depleted (Supplementary **Fig. S3I-M**). Importantly, OT-II cells primed in the Mgl2-Cre;CD11c-dlDTR mice differentiated normally into Th1 and Th2 cells under both Th2 (papain) and non-Th2 (FCA) immunization conditions (**Fig. 6H-J**), though modest reduction in their expansion was observed in both conditions (Supplementary **Fig. S3N-Q**). These results indicate that, while DCs other than CD301b^+^ DC subset may be required for the maximal OT-II cell expansion, CD301b^+^ DCs are both required and sufficient for Th2 cell fate instruction in these immunization conditions.

### CD301b^+^ DCs instruct Th2 cell fate on CD4T cells through the DC-intrinsic CD40–IL-2 axis

To further elucidate if the sufficiency of CD301b^+^ DCs for Th2 cell fate instruction is based on their cell-intrinsic CD40–IL-2 axis, we next utilized anti-CD154 (CD40L) blocking mAb (MR-1) or anti-CD40 (FGK4.5) agonistic mAb, both of which serve to inhibit physical interaction between CD40 and CD154 ^57, 58, 64, 65^. Blockade of CD154 in WT mice or genetic deficiency of CD40, but not stimulation of CD40, resulted in impaired upregulation of CD25 without affecting the CD69 expression in OT-II cells primed with OVA plus papain (**Fig. 6K-M**). Accordingly, the blockade of CD154 blunted Th2 cell differentiation of the donor OT-II cells in WT mice, whereas stimulation of CD40 restored Th2 cell differentiation even when CD154 was blocked, indicating that CD40 stimulation is sufficient to drive Th2 cell differentiation even in the absence of CD154 signaling in CD4T cells (**Fig. 6N,O**). However, the CD40 stimulation failed to restore Th2 cell differentiation when IL-2 was additionally neutralized, suggesting that IL-2 is responsible for the Th2 fate instruction at the downstream of CD40 stimulation (**Fig. 6O**). Furthermore, the CD40 stimulation could not restore Th2 cell differentiation of the donor OT-II cells in CD301b^+^ DC-depleted mice, CD301b^ΔIL-2^ mice, or in CD301b^ΔMHCII^ mice (**Fig. 6O**), indicating that the CD40 stimulation induces Th2 cell differentiation by inducing IL-2 production from the CD301b^+^ DCs that are in cognate interaction with CD4T cells. We observed no major impact on proliferation of the donor OT-II cells in any of the conditions (**Fig. 6P**). Notably, the CD40 stimulation significantly increased the frequencies of IFNγ^+^ Th1 OT-II cells in WT mice (Supplementary **Fig. S3R**). Unlike CD40-induced Th2 cell differentiation, a trend for an increase in Th1 differentiation, though statistically insignificant, was also observed in CD301b^+^ DC-depleted, CD301b^ΔIL-2^, or in CD301b^ΔMHCII^ mice upon stimulation of CD40, suggesting that the stimulation of CD40 in CD301b^−^ antigen-presenting can promote Th1 cell differentiation. Collectively, these data indicate that MHCII– and CD40-dependent interaction between CD301b^+^ DCs and antigen-specific CD4T cells and the resultant production of IL-2 from CD301b^+^ DCs play a decisive role in Th2 cell fate instruction.

### CD301b^+^ DC-intrinsic CD40–IL-2 axis skews CD4T cells toward non-Tfh effector fate

We have shown previously that the depletion of CD301b^+^ DCs in the Mgl2-DTR mice results in expansion of Tfh cells in addition to the impaired Th2 cell differentiation ^17^. This Tfh expansion is due at least partially to the loss of PD-L1-dependent suppression of Tfh cells by CD301b^+^ DCs ^17^, but it remains unclear if CD301b^+^ DCs suppress the initial fate decision of antigen-specific CD4T cells into Tfh cells. During CD4T cell priming, Tfh cells downregulate PSGL1 prior to upregulating CXCR5 ^66^. Unlike CD301b^+^ DC-depleted Mgl2-DTR mice ^17^, mature CXCR5^+^ PD-1^+^ Tfh cells were not increased in either the OT-II or endogenous CD4T cell compartment in CD301b^ΔMHCII^ mice immunized with OVA plus papain (**Fig. 7A,B**), suggesting that the expansion of mature CXCR5^+^ PD-1^+^ Tfh cells in CD301b^+^ DC-depleted mice is mainly driven by the loss of antigen-independent suppression mechanism such as PD-L1-mediated suppression of Tfh cells ^17^. However, the PSGL1^lo^ PD-1^+^ ‘immature’ Tfh fraction was significantly increased in both OT-II and endogenous CD4T cells in CD301b^ΔMHCII^ mice, suggesting that more CD4T cell are poised to the Tfh cell fate in those mice (**Fig. 7C,D**).

**Fig. 7.**
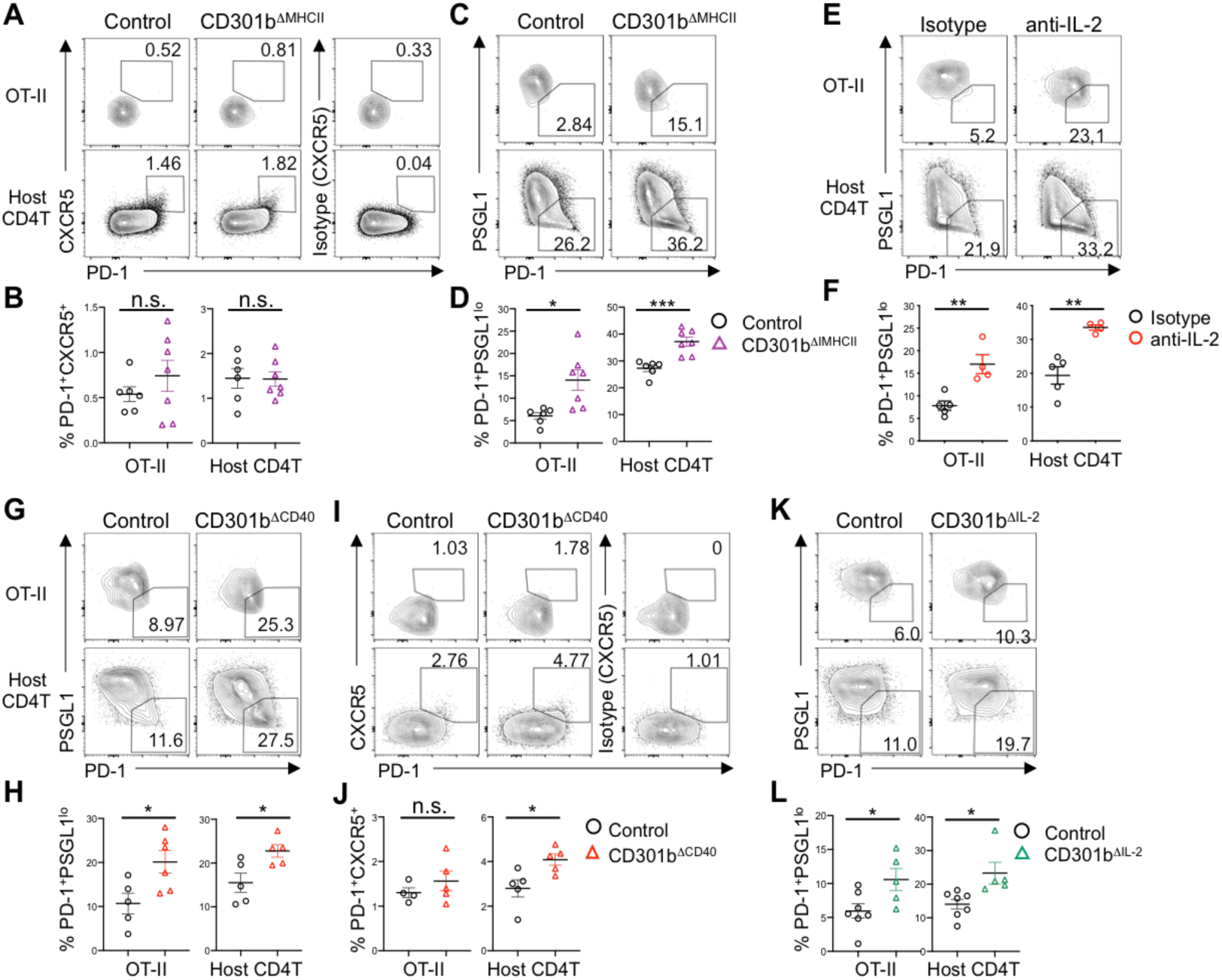
CD301b^+^ DC-intrinsic CD40–IL-2 axis skews CD4T cells toward non-Tfh effector fate. CD301b^ΔMHCII^ **(A-D)**, WT **(E, F)**, CD301b^ΔCD^^40^ **(G-J)**, CD301b^ΔIL-2^ **(K-L)**, and respective control mice were transferred with 1×10^5^ OT-II cells and immunized one day later with OVA plus papain in the footpad. In **(E, F)**, WT mice were i.p. injected with anti-IL-2 neutralizing mAb S4B6-1 or isotype control at the time of OT-II cell transfer. The dLNs were harvested on day 7 (CD301b^ΔMHCII^, WT, and CD301b^ΔCD^^40^) or day 5 (CD301b^ΔIL-2^) after the immunization. Flow cytometry plots of PD-1 and CXCR5 **(A and I)** or PD-1 and PSGL-1 **(C, E, G, K)**, and frequencies of CXCR5^+^ PD-1^+^ cells **(B, J)** or PSGL1^lo^ PD-1^+^ cells **(D, F, H, L)** among the donor CD44^+^ OT-II cells or host CD44^+^ CD4T cells are shown. Data represent means ± SEM **(B, D, F, H, J, L)**, or show representative flow cytometry plots of at least two independent experiments **(A, C, E, G, I, K)**. Not significant (n.s.), *p*≥0.05; \**p*<0.05; \*\**p*<0.01, and \*\*\**p*<0.001, by two-tailed Student’s t test **(B, D, F, H, J, L)**.

CD4T cell-intrinsic IL-2R signaling suppresses Tfh cell differentiation while favoring the differentiation of non-Tfh effector Th cells ^67, 68, 69, 70^, but the role of T cell-extrinsic source of IL-2 in this bifurcation is unknown. In WT mice transferred with OT-II cells, the blockade of IL-2 by anti-IL-2 mAb resulted in a significant increase in the differentiation of PSGL1^lo^ PD-1^+^ CD4T cells upon immunization with OVA plus papain (**Fig. 7E,F**). Notably, the differentiation of the PSGL1^lo^ PD-1^+^ CD4T cells was also increased in OT-II cells primed in CD301b^ΔCD40^ mice and CD301b^ΔIL-2^ mice compared with those primed in respective control mice, which was also observed in their endogenous CD4T cells (**Fig. 7G-L**). Interestingly, unlike in CD301b^ΔMHCII^ mice (**Fig. 7A,B**), the endogenous, but not OT-II, CXCR5^+^ PD-1^+^ mature Tfh cells were also increased in the CD301b^ΔCD40^ mice compared with the WT:Mgl2-DTR mixed BMC control mice (**Fig. 7I,J**), suggesting that the depletion of 50% of CD301b^+^ DCs in the CD301b^ΔCD40^ mice, in combination with the lack of CD40 in the remaining CD301b^+^ DCs, had more pronounced impact on Tfh cell suppression than the CD301b^+^ DC-specific MHCII deficiency alone. Thus, in addition to driving the differentiation toward Th2 cell fate, these data indicate that the CD301b^+^ DC-intrinsic CD40–IL-2 axis also limits the commitment to Tfh cell fate and promote non-Tfh effector Th cell fate (**Fig. 7G-L**). Taken together, these results demonstrate the role of CD301b^+^ DC-derived IL-2 in priming and effector fate bifurcation of CD4T cells.

## Discussion

Th2 cells play a fundamental role in type 2 adaptive immunity such as host protection against helminth parasites and pathogenesis of allergic diseases. The transcription factor GATA-3 has long been known as the cell-intrinsic master regulator for the Th2 cell identity, but the mechanism of how the Th2 cell fate is instructed by exogenous cues *in vivo* remains unclear. Although IL-4 is commonly used to induce Th2 cell differentiation in cultured CD4T cells, there is no clear evidence that DCs produce IL-4 to instruct the Th2 cell fate ^71^, and the expression of IL-4 receptor in CD4T cells is not strictly required for their Th2 fate decision *in vivo* ^72^. The present study reveals that the DC-intrinsic CD40–IL-2 axis in CD301b^+^ DCs plays a crucial role in Th2 cell fate instruction by quantitatively regulating the IL-2R signaling in CD4T cells. Upon encountering antigen-specific naive CD4T cells, CD301b^+^ DCs not only present the antigen to stimulate the TCR for timely activation but also provide IL-2 to T cells upon CD40 ligation to induce full upregulation of CD25, which is required specifically for Th2 cell differentiation, as Th2 cells rely on the IL-2R signaling for their differentiation more stringently than Th1 cells. Furthermore, the CD301b^+^ DC-derived IL-2 also limits the differentiation into Tfh cells, favoring the commitment to non-Tfh effector Th cells altogether. These data demonstrate that CD301b^+^ DC-derived IL-2 plays a decisive role in Th cell fate instruction and highlight the importance of DC-dependent quantitative control of IL-2R signaling in CD4T cells for the commitment to Th2 cell fate.

Originally identified as the T cell growth factor secreted by activated T cells themselves ^73, 74^, IL-2 plays a multifaceted role in proliferation, differentiation, and survival of CD4T cells. Although recombinant IL-2 is often added as a cell culture supplement to activated CD4T cells to maintain the survival of ‘Th0’ cells without skewing them toward the Th1 or Th2 status, previous studies have shown that CD4T cell-intrinsic production and sensing of IL-2 inhibits the differentiation of Th17 and Tfh cells ^67, 68, 69, 75, 76^, while it is required for the differentiation of Th1 and Th2 cells ^28, 29, 69, 77, 78, 79, 80^. In Th2 cell differentiation, STAT5, the main signaling component of the IL-2R, has been shown to directly bind to the *Il4*, *Il4ra* and *Gata3* loci and promote their expression ^28, 79, 80^, leading to the activation of the IL-4–IL-4Rα–STAT6 positive feedback loop to accelerate the differentiation of Th2 cells and dictate their identity ^81, 82^. Accordingly, we have shown previously that CD301b^+^ DCs are required for the upregulation of IL-4Rα in antigen-specific CD4T cells during the priming ^18^. However, STAT5 also promotes the expression of Th1-associated genes such as *Il12rb*, *Tbx21,* and *Ifng* in the context of Th1 cell differentiation ^29, 83^. Our data indicate that, while both Th1 and Th2 cell differentiation ultimately requires CD25 expression, Th2 cell differentiation is more sensitive to a partial loss of CD25. These observations suggest that quantitative differences in the IL-2R signaling during CD4T cell priming leads to qualitative differences in Th cell fate decision.

In line with the Th cell fate regulation by quantitative control of IL-2R signaling, the availability of IL-2 is tightly regulated in the LN microenvironment. Studies have shown that IL-2 produced by antigen-specific CD4T cells is rapidly quenched by Treg cells and/or DCs to prevent aberrant activation of self-reactive clones and to promote Tfh cell differentiation ^36, 37, 40, 41, 48, 50^. However, since the IL-2 in CD4T cells functions predominantly in a paracrine manner ^70^, it is unclear how the IL-2R signaling overshoots the threshold in rare antigen-specific CD4T cells immediately after priming. Our data indicate that CD301b^+^ DCs are a critical source of IL-2 early after CD4T cell priming, especially when CD4T cells require the maximal IL-2R signaling for Th2 cell differentiation. This directed action of IL-2 is facilitated by the expression of CD25 by CD301b^+^ DCs. These data suggest that IL-2 may be directly trans-presented to cognate CD4T cells through the membrane-bound CD25 as previously suggested in other contexts ^45, 51^, though the precise mechanism needs further investigation. Importantly, IL-2 production in CD301b^+^ DCs is induced by CD40, which needs to be coincided with MHCII-dependent cognate interaction for the Th2 cell fate instruction. Thus, CD301b^+^ DCs appear to ‘kick-start’ the IL-2R signaling in CD4T cells by directly handing IL-2 over to the cognate clones, thereby minimizing the IL-2 quenching effect by surrounding Treg cells. Interestingly, previous studies have shown the production of IL-2 by DCs at the interface between DCs and CD4 T cells *in vitro* ^43, 45^, suggesting that direct IL-2 delivery by CD301b^+^ DCs to CD4T cells through the immunological synapse ensures timely and confined consumption of IL-2 by the cognate clones.

While our data in CD301b^+^ DC-enriched Mgl2-cre;CD11c-dlDTR mice suggest that CD301b^+^ DCs are sufficient for inducing Th2 cell differentiation, the relatively normal Th1 cell differentiation in those mice, despite modestly impaired expansion, also indicates that the cognate interaction with CD301b^+^ DCs alone does not constrain the fate of CD4T cells to the Th2 status. Further, CD301b^+^ DCs are required for the maximal IL-2R signaling even under non-Th2 immunization condition with FCA, but, unlike Th2 cell differentiation, it seems to be dispensable for the commitment to Th1 cell fate. Notably, in CD4T cells stimulated with cognate peptides *in vitro*, the dose of the peptide has been shown to correlate with the Th1-to-Th2 ratio while inversely correlating with STAT5 activation due to inhibition of STAT5 by ERK, a signal downstream of the TCR, implicating that Th1 cells only require a lower amount of activated STAT5 for their differentiation while Th2 cell differentiation requires more potent STAT5 activation ^28^. Moreover, a recent study found that IL-6 induced by antigen-independent inflammation dampens Th2 and promote Th17 cell differentiation by prematurely down-regulating CD25 expression in activated CD4T cells ^84^. Thus, the non-Th2 fate appears to be associated with excess TCR signaling and/or inflammation that results in a reduced IL-2R signaling. We reported recently that CD301b^+^ DCs in the LNs are located in closer proximity to high endothelial venules and have earlier access to incoming naive CD4T cells than LCs and cDC1s, which makes them critical for timely priming of antigen-specific CD4T cells ^18^. Given that LCs and cDC1s encounter CD4T cells in a delayed timing from CD301b^+^ DCs but are often required for non-Th2 Th cell differentiation ^3, 4, 85, 86^, under non-Th2 immunization conditions, the Th2-biased status imprinted early after priming by CD301b^+^ DCs may be overridden by the non-Th2 fate through the subsequent interaction with other DCs, as we recently discussed elsewhere ^87^. The nature of such staggered interaction with multiple DC subsets and the role of CD301b^+^ DCs in these conditions need further investigation, as the non-Th2 fate instruction cues may be delivered through antigen-independent interactions ^84, 85^.

Taken together, our data demonstrate that CD301b^+^ DC-intrinsic CD40–IL-2 axis drives Th2 cell fate instruction by quantitatively regulating the IL-2R signaling in antigen-specific CD4T cells. While the use of a simplistic immunization approach in this study allows precise analysis of the T cell priming kinetics, the mechanism for disease-associated Th2 cell differentiation may need to be examined in a more complex setting such as allergies and helminth infection. In particular, whether the excess amount of type 2 cytokines released from innate immune cells under these conditions overrides the requirement of DC-derived IL-2 at the time of priming needs to be addressed in future studies.

## Methods

### Mice

Mgl2-DTR and Mgl2-Cre mice were a gift from Akiko Iwasaki (Yale University). *Il2^flox/flox^* mice ^47^ were provided by Weizhou Zhang (University of Florida). OT-II mice on the IL-4-GFP background (4get;OT-II) were a gift from Jason Weinstein (Rutgers New Jersey Medical School). Generation of Mgl2-DTR (*Mgl2^+/DTR-EGFP^*) ^11^, Mgl2-Cre and CD11c-dlDTR mice ^18^ were previously described and maintained in our colony. C57BL/6N (B6) and congenic CD45.1 (B6.SJL-*Ptprc^a^Pepc^b^*/BoyCrCrl) mice on B6 background were purchased from Charles River Laboratory and propagated in our colony. *H2-Ab1* (MHCII) ^flox/flox^ (B6.129X1-*H2-Ab1^b-tmKoni^*/J), *Il2ra^flox/flox^* (B6(129S4)-*Il2ra^tm1c^*(EUCOMM)*^Wts^*^i^/TrmaJ), CD40^−/−^ (B6.129P2-*Cd40^tm1Kik^*/J), MHCII^−/−^ (B6.129S2-*H2^dlAb^*^1^*^-Ea^*/J), OT-II (B6.Cg-Tg(TcraTcrb)425Cbn/J), CD207-DTR (B6.129S2-*Cd207^tm^*^3^.^1^(HBEGF/EGFP)*^Ma^*^l/^J), *Rag1^−/−^* (B6.129S7-*Rag1^tm1Mom^*/J), Il2ra^−/−^ (B6;129S4-*Il2ra^tm1Dw^*/J), Rosa26 loxP-stop-loxP (LSL)-tdTomato (Ai14, R26^LSL-iTom^) reporter (B6.Cg-*Gt(ROSA)26Sor^tm^*^14^*^(CAG-tdTomato)Hze^*/J), and Nur77-GFP reporter (C57BL/6-Tg(Nr4a1-EGFP/cre)820Khog/J) mice were obtained from the Jackson Laboratory (Bar Harbor, ME) and maintained in-house.

*Mgl2^cre/cre^* mice were crossed with *H2-Ab1*^flox/flox^, *Il2^flox/flox^*, and Il2ra^flox/flox^ mice to generate *Mgl2^+/cre^*;*H2-* ^Ab1flox/flox^ (CD301b^ΔMHCII^), Mgl2*^+/cre^*;Il2ra*^flox/flox^* (CD301b^ΔIL-2^), and *Mgl2*^+/cre^;Il2ra^flox/flox^ (CD301b^ΔCD25^) mice, respectively. The leaky expression of Cre in the *Mgl2^+/cre^* mice were monitored by flow-cytometry or by qPCR and those with global deletion of MHCII, CD25 or IL-2 were excluded from the analysis as previously described ^18^. CD11c (*Itgax*)-dlDTR mice were crossed with Mgl2-Cre mice to generate *Mgl2^+/Cre^*;*Itgax^+/dlDTR^*mice. The *Mgl2^+/Cre^*;*Itgax^+/dlDTR^* mice were further crossed with R26^LSL-iTom^ reporter mice to produce *Mgl2^+/Cre^*;*Itgax^+/dlDTR^*;*Rosa26^+/LSL-iTom^*mice. OT-II and 4get;OT-II mice were bred to CD45.1 mice to produce CD45.1;OT-II and CD45.1;4get;OT-II mice, respectively. Nur77-GFP mice were crossed with CD45.1;OT-II mice to generate CD45.1;Nur77-GFP;OT-II mice. *Il2ra^−/−^* mice were crossed with CD45.1;OT-II mice to generate CD45.1;*Il2ra^+/−^*;OT-II and CD45.1;I*l2ra^−/−^*;OT-II mice. The CD45.1;*Il2ra^+/−^*;OT-II and CD45.1;*Il2ra^−/−^*;OT-II mice were maintained on the *Rag1^−/−^*background and further crossed onto the Nur77-GFP background where indicated. All mice were maintained in a specific pathogen-free facility at Rutgers New Jersey Medical School. All animal experiments in this study have been approved by the Institutional Animal Care and Use Committee at Rutgers University.

### BM reconstitution

For generating CD40*^−/−^*;Mgl2-DTR (CD301b^ΔCD40^) and CD40*^−/−^*;MHCII*^−/−^*mixed BMC mice, lethally irradiated (1100 cGy) B6 mice were reconstituted with a 50:50 mixture of CD40*^−/−^* and Mgl2-DTR BM cells, or with CD40*^−/−^* and MHCII*^−/−^* BM cells (2×10^6^ cells each), respectively. Those reconstituted with WT and CD40*^−/−^* BM cells were used as a control for CD301b^ΔCD40^ and CD40*^−/−^*;MHCII*^−/−^* mixed BMC mice. BMC mice were maintained on antibiotics (sulfamethoxazole and trimethoprim, NDC 0121-0854-16, PAI pharma) in their drinking water for 2 weeks after the reconstitution and rested for at least 6-8 weeks before being used for experiments.

### Immunization and *in vivo* treatment

All immunization procedures were conducted in the rear footpad with 20 µL injection volume per footpad. Mice were immunized as indicated in each Figure with 5 µg low-endotoxin OVA (Worthington Biochemical Corporation) mixed with either 50 µg papain (P4762, Sigma), 10 µL FCA (F5881, Sigma), or 10 µL alum (77161, Thermo Scientific) in phosphate-buffered saline (PBS). For DC depletion, mice were injected i.p. with 500 ng DT in PBS (List Biological Laboratories) at indicated time-points. For CD40 stimulation of DCs *in vivo*, mice were injected i.p. with 100 µg agonistic anti-CD40 mAb (FGK4.5, BioXcell) in PBS. LNs were harvested 24 hours later and restimulated with PMA and ionomycin for intracellularly staining IL-2. For mAb treatment of mice immunized with OVA plus papain, mice were injected i.p. with anti-IL-2 (S4B6-1, BioXcell, 100 µg), anti-CD154 (MR1, BioXcell, 100 µg), anti-CD40 (FGK4.5, BioXcell, 100 µg), anti-CD154 plus anti-CD40 (100 µg each), or anti-IL-2 plus anti-CD154 plus anti-CD40 (100 µg each) mAbs at the time of OT-II transfer or immunization as indicated. Rat IgG2a isotype control (2A3, BioXcell) mAb was injected to make the amount of mAbs equal across different groups.

### T cell priming assays

For adoptively transferring OT-II cells, CD4 T cells were isolated from the spleen and LNs of indicated naïve OT-II strains with Mouse CD4 T Cell Isolation Kit (STEMCELL Technologies 19852 or BioLegend 480033) and labeled with 1.0 µM CFSE (Thermo Fisher) or CTV (C34571, Thermo Fisher) according to the manufacturer’s protocol. The purity of isolated OT-II cells was typically 90-95%. For analyzing the CD4 T cell activation kinetics, CFSE– or CTV-labelled CD45.1;OT-II cells (1 to 2×10^6^) were adoptively transferred into CD45.2/.2 recipient mice that had been immunized in the rear right footpad 24 hours prior. Mice with DTR expression were treated with DT as indicated in each Figure. In some experiments, multiple types of donor cells (1×10^6^ cells each) were co-transferred into one mouse. Two hours after the OT-II cell transfer, further LN entry of T cells was blocked by retro-orbitally injecting 100 µg anti-CD62L mAb (clone Mel-14, BE0021, BioXCell). The left (ndLN) and right (dLN) popliteal LNs were harvested at indicated time-points. For analyzing CD4T cell differentiation, CFSE– or CTV-labeled OT-II cells (1×10^5^ cells) were adoptively transferred into indicated CD45.2/.2 recipient mice, followed by immunization with OVA and an indicated adjuvant in the footpad. The dLNs were harvested at indicated time-points for intracellularly staining cytokines.

### Cell preparations and flow cytometry

For preparing single cell suspensions, LNs were minced and enzymatically digested with 2.5 mg/ml collagenase D (11088882001, Sigma) in complete RPMI-1640 medium with 10% heat-inactivated fetal bovine serum (FBS) at 37°C for 30 min. The cells were washed with PBS containing 2mM EDTA, and then stained for dead cells with cell viability dye (Zombie Aqua or Zombie UV, BioLegend) in PBS on ice for 20 min. The cells were washed with 2mM EDTA/PBS, incubated with 10 µg/mL anti-CD16/CD32 (2.4G2, BioLegend) on ice for 10 min to block non-specific antibody binding, and stained with fluorochrome-conjugated mAbs on ice for 20 min. For intracellular cytokine staining, LN single cell suspensions were stimulated in a 96-well round-bottom plate with Cell Stimulation Cocktail containing PMA and ionomycin (eBioscience 00-4970-03, Thermo Fisher or 423302, BioLegend) at 37 °C for 1 hour, and then incubated for another 5 hours at 37 °C with additional Protein Transport Inhibitor Cocktail containing Brefeldin A and Monensin (eBioscience 00-4980-03, Thermo Fisher). Cells were then fixed and permeabilized with BD Cytofix/Cytoperm Kit (BD Biosciences) and incubated with anti-cytokine mAbs for 30 min on ice. For staining phosphorylated STAT5 and GATA-3, LNs were fixed with BD Phosflow^TM^ Fix Buffer I (BD Biosciences) at 37 °C for 10 min immediately after the harvest and permeabilized with BD Phosflow^TM^ Perm Buffer III (BD Biosciences) on ice for 30 min. After blocking with anti-CD16/CD32 (2.4G2, BioLegend), cells were stained for cell surface molecules on ice for 20 min, washed with PBS containing 2 mM EDTA, and then incubated with anti-pSTAT5 and anti-GATA-3 mAbs at room temperature (20°C–25°C) for 16 hours.

The mAbs to CD4 (RM4-5 or GK1.5), CD8α (53-6.7), CD45R/B220 (RA3-6B2), TCRβ (H57-597), Ly6G (1A8), CD69 (H1.2F3), CD25 (PC61), CD44 (IM7), CD279 (PD-1) (RMP1-30), I-A/I-E (MHCII) (M5/114.15.2), CD11b (M1/70), CD11c (N418), CD326 (G8.8), CD301b (URA1), CD172a (P84), CD273 (PD-L2) (TY25), XCR1 (ZET), CD103 (2E7), CD40 (3/23), CD80 (16-10A1), CD86 (GL-1), CD45.1 (A20), CD45.2 (104), IFNγ (XMG1.2), IL-4 (11B11), IL-2 (JES6-5H4), and GATA-3 (16E10A23) were purchased from BioLegend. Anti-pSTAT5 (Tyr694) mAb (SRBCZX) was purchased from eBioscience. Anti-CXCR5 (2G8) and PSGL-1 (2PH1) mAbs were purchased from BD Biosciences. All mAbs used for flow cytometry were prepared in FACS buffer (1% BSA, 2mM EDTA, 0.05% sodium azide in PBS), except that mAbs to pSTAT5 and GATA-3 were prepared in 1× BD Perm/Wash buffer (BD Biosciences).

All samples were collected on BD LSRII (BD Biosciences) or Attune NxT (Thermo Fisher) flow cytometer and analyzed with FlowJo software (Version 9.3.2 and 10.5.0, BD).

### CITEseq data acquisition and analysis

For CITEseq, B6 mice were immunized with 5 ug OVA plus 50 ug papain in both rear footpads in 20 µL PBS. Popliteal lymph nodes were harvested from immunized mice 1 day later and digested with collagenase D (2.5 mg/mL, Roche) in RPMI-1640 medium supplemented with 10% FBS for 30 min at 37°C, after which EDTA (5mM final conc.) was added and incubated for another 10 min to allow complete dissociation. Pooled skin-dLNs from naive B6 mice were used as a control. Single cell suspensions were collected, and cells were stained with viability dye (Zombie Aqua, BioBegend) in PBS for 20 minutes on ice. Cells were washed and incubated with anti-CD16/32 (10ug/mL, clone 2.4G2, BioLegend) for 20 minutes on ice and then stained with fluorochrome-labeled mAbs for B220 and MHCII and DNA-barcoded TotalSeq mAbs for MHCII, CD11c, CD301b, CD8α, CD11b, CD64, CD103, CD172a, Ly6C and XCR1 (BioLegend) in PBS containing 1%BSA and 2mM EDTA for 20 min on ice. The TotalSeq mAbs for MHCII, CD11c and CD301b (0.25 mg/mL) were pre-mixed with AlexaFluor 488-, AlexaFluor 546, and AlexaFluor 647-labeled antisense oligoniculeotide (1 µM) that is specific to each DNA barcode, respectively, for 15 minutes at room temperature (20°C–25°C) in order to detect these antibodies by flow cytometry for sorting. Live MHCII^hi^ CD11c^+^ B220^−^ cells were sorted by FACS Aria III cell sorter (BD) and used to generate Gel Beads-in-Emulsion (GEMs) using the Chromium Single-Cell 3’ Reagent v3 kit (1000092, 10X Genomics) according to the manufacturer’s protocol. Briefly, cell suspensions with a >90% viability was mixed with reverse transcription reagents and loaded onto Chromium Chip B (1000074, 10X Genomics) along with Gel beads and Partitioning oil in the recommended order, and then the chip was processed through 10X Chromium Controller for the generation of GEMs, followed by reverse transcription, cleanup, and cDNA amplification. After the cDNA amplification, size selection was used to separate the cDNA molecules for 3’ Gexp (Gene Expression) library and ADT library construction. Library quantity and quality control were performed on Qubit 4 Fluorometer using HS reagent kit, (Q33231, Invitrogen) and TapeStation using HS DNA D1000 screen tape (5067-5584, Agilent Technologies). ADT and 3’ Gexp libraries were mixed at the ratio of 1:5 and sequenced on NovaSeq 6000 sequencer (Illumina) with a configuration of 28/8/0/91-bp for cell barcode, sample barcode and mRNA reads, respectively, as recommended by 10X Genomics. Cell Ranger and Loupe browser were used for data analysis. The aligned data were further analyzed and visualized using Partek Flow (Partek Inc) and Ingenuity Pathway Analysis (QIAGEN) software.

## Statistical analysis

P-values were calculated by two-tailed Student’s unpaired t test or by paired t test as indicated using Prism software (version 8, GraphPad). Differences were defined as statistically significant when P-values were < 0.05. Data are presented as mean ± SEM.

## Data availability

The CITEseq data used in this study will be deposited in the Gene Expression Omnibus (GEO) database at NCBI upon formal publication of this manuscript. All other data and materials in this study are available from the corresponding author upon request with an appropriate material transfer agreement.

## Acknowledgements

We thank Kendall A. Smith for sharing the *Il2^flox/flox^*mice. We thank Weizhou Zhang and Akiko Iwasaki for the original breeders of the *Il2^flox/flox^*mice and Mgl2-Cre and Mgl2-DTR mice, respectively. We thank George Yap for critical reading of the manuscript. We thank the Genomics Center, Comparative Medicine Resources, and the Flow Cytometry Core at Rutgers New Jersey Medical School for technical assistance. This work was supported by NIH grants R01AI132576, R01AI165622 and R21CA259541 to Y.K. N.T. and Y.K. designed experiments. N.T. performed most of the experiments with help from J.E.-F. and A.D.-P. J.E.-F. and Y.K. performed the CITEseq experiments and analyzed the data with assistance from the Flow Cytometry Core and the Genomics Center at Rutgers New Jersey Medical School. N.T., J.E.-F. and Y.K. performed statistical and computational analyses and interpreted the data. Y.K. conceptualized and supervised the study and acquired funding. N.T. and Y.K. wrote the paper. The authors declare that they have no competing interests. The CITEseq data used in this study will be deposited in GEO database upon formal publication. Mgl2-DTR and Mgl2-Cre mice have been deposited in the Jackson Laboratory. CD11-dlDTR mice are available upon request from Y.K. under an MTA from Rutgers University.

---

**Fig. S1.**
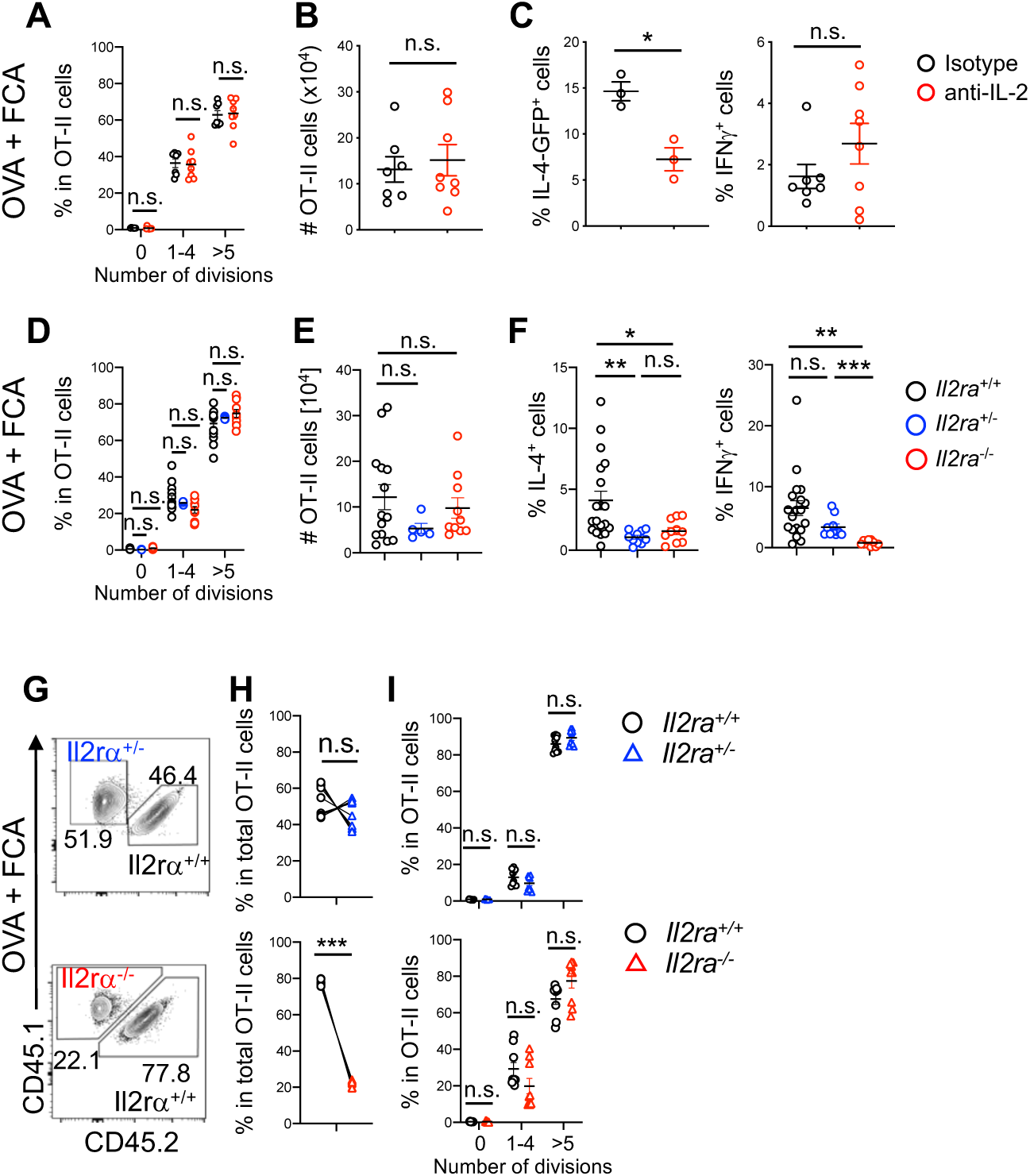
Effect of reduced IL-2R signaling on CD4T cell differentiation under a non-Th2 immunization condition with FCA. **A-C**. As in Fig. 3A, but the mice were transferred with 4get;OT-II cells and immunized with OVA plus FCA. Frequencies of OT-II cells that have undergone indicated number of cell divisions **(A)**, numbers of OT-II cells **(B)**, and frequencies of IL-4^+^ and IFNγ^+^ cells among the donor OT-II cells **(C)** are shown. **D-F.** As in Fig. 3K, but the mice were immunized with OVA plus FCA. Frequencies of OT-II cells that have undergone indicated number of cell divisions **(D)**, numbers of OT-II cells **(E)**, and frequencies of IL-4^+^ and IFNγ^+^ cells among the donor OT-II cells **(F)** are shown. **G-I.** As in Fig. 3R, but the mice were immunized with OVA plus FCA. Representative flow cytometry plots of CD45.1/.1 (*Il2ra^+/−^*or *Il2ra^−/−^*) and CD45.1/.2 (*Il2ra^+/+^*) OT-II cells among the total CD45.1^+^ donor OT-II cells **(G)**, paired frequencies of indicated OT-II cells among total OT-II cells within individual host dLNs **(H)**, and frequencies of indicated OT-II cells that have undergone indicated number of cell divisions **(I)** are shown. In **(H)**, each connecting line represents OT-II cells of two different genotypes isolated from the same host. Data represent means ± SEM **(A-F, H-I)**, or show representative flow cytometry plots of at least two independent experiments **(G)**. Not significant (n.s.), *p*≥0.05; \**p*<0.05; \*\**p*<0.01; and \*\*\**p*<0.001, by two-tailed Student’s t test **(A-F, I)** or paired t test **(H)**.

**Fig. S2.**
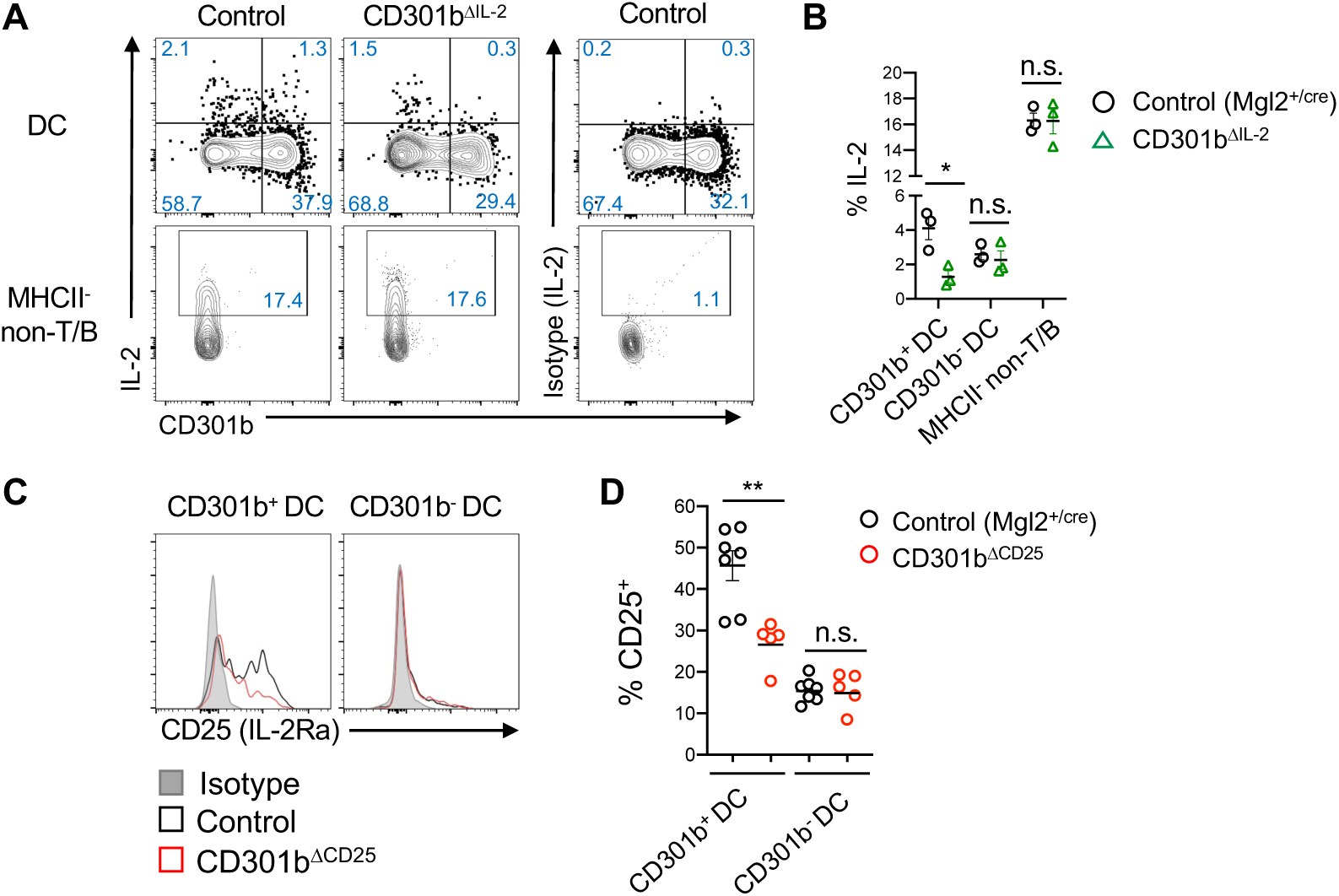
Specific deletion of IL-2 and CD25 in CD301b^ΔIL-2^ and CD301b^ΔCD25^ mice. **A-B**. Control (Mgl2^+/Cre^) and CD301b^ΔIL-2^ mice were immunized with papain in the footpad. The dLNs were harvested 24 hours after the immunization and restimulated *ex vivo* with PMA and ionomycin for intracellularly staining IL-2. Flow cytometry plots **(A)** and frequencies **(B)** of IL-2 in CD301b^+^ DCs, CD301b^−^ DCs, or MHCII^−^ non-T/B cells as gated in Fig. 4B are shown. **C-D**. CD25 expression was examined in the skin-dLNs of naïve control (*Mgl2^+/Cre^*) and CD301b^ΔCD^^25^ mice. Flow cytometry histograms **(C)** and frequencies **(D)** of CD25 in CD301b^+^ and CD301b^−^ DCs are shown. Data show representative flow cytometry plots of at least two independent experiments **(A, C)** or represent means ± SEM **(B, D)**. Not significant (n.s.), *p*≥0.05; \**p*<0.05; and \*\**p*<0.01, by two-tailed Student’s t test **(B, D)**.

**Fig. S3.**
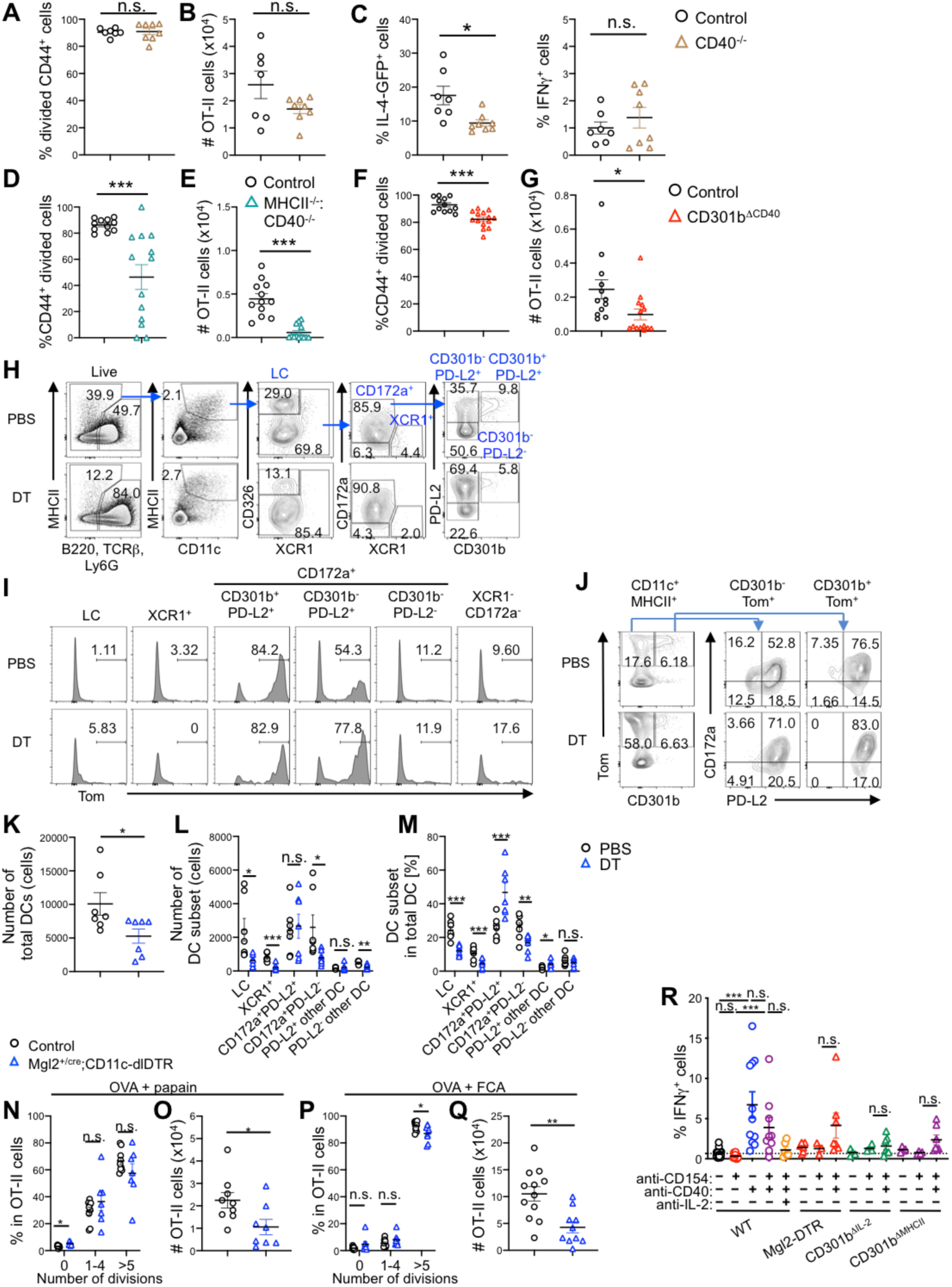
CD301b^+^ DC-intrinsic CD40–IL-2 axis is necessary and sufficient for Th2 cell fate instruction. **A-C**. WT and CD40^−/−^ mice were transferred with 1×10^5^ CTV-labeled 4get;OT-II cells and immunized one day later with OVA plus papain in the footpad. The dLNs were harvested 5 days after the immunization. Frequencies of CD44^+^ CFSE^lo^ divided cells among the donor OT-II cells **(A)**, numbers of OT-II cells **(B)**, and frequencies of IL-4-GFP^+^ and IFNγ^+^ cells among the donor OT-II cells **(C)** are shown. **D-G.** Frequencies of CD44^+^ CFSE^lo^ divided cells among the donor OT-II cells **(D, F)** and numbers of OT-II cells **(E, G)** in the dLNs of mice shown in Fig. 6A **(D, E)** and Fig. 6D **(F, G)** are shown. **H-M.** Mgl2^+/cre^;CD11c-dlDTR;R26^LSL-iTom^ mice were treated with PBS or DT, and inguinal LNs were harvested 2 days after the DT treatment and stained for DC subsets. Gating strategy for DC subsets including LCs, XCR1^+^ CD172^−^ cDC1s, CD301b^+^ PD-L2^+^ CD172a^+^ cDC2s, CD301b^−^ PD-L2^+^ CD172a^+^ cDC2s, CD301b^−^ PD-L2^−^ CD172a^+^ cDC2s, and other DCs (XCR1^−^ CD172a^−^) **(H)**, representative flow cytometry histograms of tdTomato expression in each DC subset **(I)**, representative flow cytometry plots of tdTomato and CD301b expression on total DCs (CD11c^+^MHCII^+^) and the expression of CD172a and PD-L2 on tdTomato^+^ CD301b^+^ and tdTomato^+^ CD301b^−^ populations **(J)**, numbers of total DCs **(K)** and each DC subset **(L)**, and frequencies of each subset among total DCs **(M)** in the inguinal LNs 2 days after DT treatment are shown. **N-Q.** Frequencies of OT-II cells that have undergone indicated number of cell divisions **(N, P)**, numbers of OT-II cells **(O, Q)** in the dLNs of mice shown in Fig. 6I **(N, O)** and Fig. 6J **(P, Q)** are shown. **R.** Frequencies of IFNγ^+^ cells among the donor OT-II cells in the dLN of mice shown in **Fig. 6N-P** are shown. Data represent means ± SEM (**A-G, K-R)**, or show representative flow cytometry plots of at least two independent experiments **(H-J)**. Not significant (n.s.), *p*≥0.05; \**p*<0.05; \*\**p*< 0.01; and \*\*\**p*<0.001, by two-tailed Student’s t test **(A-G, K-R)**.

## References

1. Zhu, J., Yamane, H. & Paul, W.E. Differentiation of effector CD4 T cell populations (*). Annu Rev Immunol 28, 445–489 (2010).

2. Yin, X., Chen, S. & Eisenbarth, S.C. Dendritic Cell Regulation of T Helper Cells. Annu Rev Immunol 39, 759–790 (2021).

3. Igyarto, B.Z. et al. Skin-resident murine dendritic cell subsets promote distinct and opposing antigen-specific T helper cell responses. Immunity 35, 260–272 (2011).

4. Martinez-Lopez, M., Iborra, S., Conde-Garrosa, R. & Sancho, D. Batf3-dependent CD103+ dendritic cells are major producers of IL-12 that drive local Th1 immunity against Leishmania major infection in mice. Eur J Immunol 45, 119–129 (2015).

5. Kashem, S.W. et al. Candida albicans morphology and dendritic cell subsets determine T helper cell differentiation. Immunity 42, 356–366 (2015).

6. Lewis, K.L. et al. Notch2 receptor signaling controls functional differentiation of dendritic cells in the spleen and intestine. Immunity 35, 780–791 (2011).

7. Persson, E.K. et al. IRF4 transcription-factor-dependent CD103(+)CD11b(+) dendritic cells drive mucosal T helper 17 cell differentiation. Immunity 38, 958–969 (2013).

8. Satpathy, A.T. et al. Notch2-dependent classical dendritic cells orchestrate intestinal immunity to attaching-and-effacing bacterial pathogens. Nat Immunol 14, 937–948 (2013).

9. Schlitzer, A. et al. IRF4 transcription factor-dependent CD11b+ dendritic cells in human and mouse control mucosal IL-17 cytokine responses. Immunity 38, 970–983 (2013).

10. Gao, Y. et al. Control of T helper 2 responses by transcription factor IRF4-dependent dendritic cells. Immunity 39, 722–732 (2013).

11. Kumamoto, Y. et al. CD301b(+) dermal dendritic cells drive T helper 2 cell-mediated immunity. Immunity 39, 733–743 (2013).

12. Leon, B. et al. Regulation of T(H)2 development by CXCR5+ dendritic cells and lymphotoxin-expressing B cells. Nat Immunol 13, 681–690 (2012).

13. Mayer, J.U. et al. Homeostatic IL-13 in healthy skin directs dendritic cell differentiation to promote T(H)2 and inhibit T(H)17 cell polarization. Nat Immunol 22, 1538–1550 (2021).

14. Tussiwand, R. et al. Klf4 expression in conventional dendritic cells is required for T helper 2 cell responses. Immunity 42, 916–928 (2015).

15. Murakami, R. et al. A unique dermal dendritic cell subset that skews the immune response toward Th2. PLoS One 8, e73270 (2013).

16. Sokol, C.L., Camire, R.B., Jones, M.C. & Luster, A.D. The Chemokine Receptor CCR8 Promotes the Migration of Dendritic Cells into the Lymph Node Parenchyma to Initiate the Allergic Immune Response. Immunity 49, 449–463 e446 (2018).

17. Kumamoto, Y., Hirai, T., Wong, P.W., Kaplan, D.H. & Iwasaki, A. CD301b+ dendritic cells suppress T follicular helper cells and antibody responses to protein antigens. Elife 5 (2016).

18. Tatsumi, N., Codrington, A.L., El-Fenej, J., Phondge, V. & Kumamoto, Y. Effective CD4 T cell priming requires repertoire scanning by CD301b(+) migratory cDC2 cells upon lymph node entry. Sci Immunol 6, eabg0336 (2021).

19. Barnden, M.J., Allison, J., Heath, W.R. & Carbone, F.R. Defective TCR expression in transgenic mice constructed using cDNA-based alpha– and beta-chain genes under the control of heterologous regulatory elements. Immunol Cell Biol 76, 34–40 (1998).

20. Bursch, L.S. et al. Identification of a novel population of Langerin+ dendritic cells. J Exp Med 204, 3147–3156 (2007).

21. Ginhoux, F. et al. Blood-derived dermal langerin+ dendritic cells survey the skin in the steady state. J Exp Med 204, 3133–3146 (2007).

22. Poulin, L.F. et al. The dermis contains langerin+ dendritic cells that develop and function independently of epidermal Langerhans cells. J Exp Med 204, 3119–3131 (2007).

23. Kissenpfennig, A. et al. Dynamics and function of Langerhans cells in vivo: dermal dendritic cells colonize lymph node areas distinct from slower migrating Langerhans cells. Immunity 22, 643–654 (2005).

24. Moran, A.E. et al. T cell receptor signal strength in Treg and iNKT cell development demonstrated by a novel fluorescent reporter mouse. J Exp Med 208, 1279–1289 (2011).

25. Osum, K.C. & Jenkins, M.K. Toward a general model of CD4(+) T cell subset specification and memory cell formation. Immunity 56, 475–484 (2023).

26. Spolski, R., Li, P. & Leonard, W.J. Biology and regulation of IL-2: from molecular mechanisms to human therapy. Nat Rev Immunol 18, 648–659 (2018).

27. Hwang, E.S., White, I.A. & Ho, I.C. An IL-4-independent and CD25-mediated function of c-maf in promoting the production of Th2 cytokines. Proc Natl Acad Sci U S A 99, 13026–13030 (2002).

28. Yamane, H., Zhu, J. & Paul, W.E. Independent roles for IL-2 and GATA-3 in stimulating naive CD4+ T cells to generate a Th2-inducing cytokine environment. J Exp Med 202, 793–804 (2005).

29. Liao, W., Lin, J.X., Wang, L., Li, P. & Leonard, W.J. Modulation of cytokine receptors by IL-2 broadly regulates differentiation into helper T cell lineages. Nat Immunol 12, 551–559 (2011).

30. Van Parijs, L. et al. Functional responses and apoptosis of CD25 (IL-2R alpha)-deficient T cells expressing a transgenic antigen receptor. J Immunol 158, 3738–3745 (1997).

31. Le Saout, C. et al. IL-2 mediates CD4+ T cell help in the breakdown of memory-like CD8+ T cell tolerance under lymphopenic conditions. PLoS One 5, e12659 (2010).

32. Setoguchi, R., Hori, S., Takahashi, T. & Sakaguchi, S. Homeostatic maintenance of natural Foxp3(+) CD25(+) CD4(+) regulatory T cells by interleukin (IL)-2 and induction of autoimmune disease by IL-2 neutralization. J Exp Med 201, 723–735 (2005).

33. Boyman, O., Kovar, M., Rubinstein, M.P., Surh, C.D. & Sprent, J. Selective stimulation of T cell subsets with antibody-cytokine immune complexes. Science 311, 1924–1927 (2006).

34. Gratz, I.K. et al. Cutting Edge: memory regulatory t cells require IL-7 and not IL-2 for their maintenance in peripheral tissues. J Immunol 190, 4483–4487 (2013).

35. Villarino, A.V. et al. Helper T cell IL-2 production is limited by negative feedback and STAT-dependent cytokine signals. J Exp Med 204, 65–71 (2007).

36. Liu, Z. et al. Immune homeostasis enforced by co-localized effector and regulatory T cells. Nature 528, 225–230 (2015).

37. Wong, H.S. et al. A local regulatory T cell feedback circuit maintains immune homeostasis by pruning self-activated T cells. Cell 184, 3981–3997 e3922 (2021).

38. Pimentel-Muinos, F.X., Munoz-Fernandez, M.A. & Fresno, M. Control of T lymphocyte activation and IL-2 receptor expression by endogenously secreted lymphokines. J Immunol 152, 5714–5722 (1994).

39. Waysbort, N., Russ, D., Chain, B.M. & Friedman, N. Coupled IL-2-dependent extracellular feedbacks govern two distinct consecutive phases of CD4 T cell activation. J Immunol 191, 5822–5830 (2013).

40. Busse, D. et al. Competing feedback loops shape IL-2 signaling between helper and regulatory T lymphocytes in cellular microenvironments. Proc Natl Acad Sci U S A 107, 3058–3063 (2010).

41. Feinerman, O. et al. Single-cell quantification of IL-2 response by effector and regulatory T cells reveals critical plasticity in immune response. Mol Syst Biol 6, 437 (2010).

42. Granucci, F. et al. Inducible IL-2 production by dendritic cells revealed by global gene expression analysis. Nat Immunol 2, 882–888 (2001).

43. Granucci, F., Feau, S., Angeli, V., Trottein, F. & Ricciardi-Castagnoli, P. Early IL-2 production by mouse dendritic cells is the result of microbial-induced priming. J Immunol 170, 5075–5081 (2003).

44. Mencarelli, A. et al. Calcineurin-mediated IL-2 production by CD11c(high)MHCII(+) myeloid cells is crucial for intestinal immune homeostasis. Nat Commun 9, 1102 (2018).

45. Wuest, S.C. et al. A role for interleukin-2 trans-presentation in dendritic cell-mediated T cell activation in humans, as revealed by daclizumab therapy. Nat Med 17, 604–609 (2011).

46. Zelante, T. et al. CD103(+) Dendritic Cells Control Th17 Cell Function in the Lung. Cell Rep 12, 1789–1801 (2015).

47. Popmihajlov, Z., Xu, D., Morgan, H., Milligan, Z. & Smith, K.A. Conditional IL-2 Gene Deletion: Consequences for T Cell Proliferation. Front Immunol 3, 102 (2012).

48. Leon, B., Bradley, J.E., Lund, F.E., Randall, T.D. & Ballesteros-Tato, A. FoxP3+ regulatory T cells promote influenza-specific Tfh responses by controlling IL-2 availability. Nat Commun 5, 3495 (2014).

49. Kim, M., Kim, T.J., Kim, H.M., Doh, J. & Lee, K.M. Multi-cellular natural killer (NK) cell clusters enhance NK cell activation through localizing IL-2 within the cluster. Sci Rep 7, 40623 (2017).

50. Li, J., Lu, E., Yi, T. & Cyster, J.G. EBI2 augments Tfh cell fate by promoting interaction with IL-2-quenching dendritic cells. Nature 533, 110–114 (2016).

51. Jones, M.C. et al. CD4 Effector TCR Avidity for Peptide on APC Determines the Level of Memory Generated. J Immunol 210, 1950–1961 (2023).

52. Fan, M.Y. et al. Differential Roles of IL-2 Signaling in Developing versus Mature Tregs. Cell Rep 25, 1204–1213 e1204 (2018).

53. Toomer, K.H. et al. Essential and non-overlapping IL-2Ralpha-dependent processes for thymic development and peripheral homeostasis of regulatory T cells. Nat Commun 10, 1037 (2019).

54. Stoeckius, M. et al. Simultaneous epitope and transcriptome measurement in single cells. Nat Methods 14, 865–868 (2017).

55. Kramer, A., Green, J., Pollard, J., Jr. & Tugendreich, S. Causal analysis approaches in Ingenuity Pathway Analysis. Bioinformatics 30, 523–530 (2014).

56. Bennett, S.R. et al. Help for cytotoxic-T-cell responses is mediated by CD40 signalling. Nature 393, 478–480 (1998).

57. Rolink, A., Melchers, F. & Andersson, J. The SCID but not the RAG-2 gene product is required for S mu-S epsilon heavy chain class switching. Immunity 5, 319–330 (1996).

58. Schoenberger, S.P., Toes, R.E., van der Voort, E.I., Offringa, R. & Melief, C.J. T-cell help for cytotoxic T lymphocytes is mediated by CD40-CD40L interactions. Nature 393, 480–483 (1998).

59. Jenkins, S.J., Perona-Wright, G. & MacDonald, A.S. Full development of Th2 immunity requires both innate and adaptive sources of CD154. J Immunol 180, 8083–8092 (2008).

60. MacDonald, A.S., Straw, A.D., Dalton, N.M. & Pearce, E.J. Cutting edge: Th2 response induction by dendritic cells: a role for CD40. J Immunol 168, 537–540 (2002).

61. Mohrs, M., Shinkai, K., Mohrs, K. & Locksley, R.M. Analysis of type 2 immunity in vivo with a bicistronic IL-4 reporter. Immunity 15, 303–311 (2001).

62. Allenspach, E.J., Lemos, M.P., Porrett, P.M., Turka, L.A. & Laufer, T.M. Migratory and lymphoid-resident dendritic cells cooperate to efficiently prime naive CD4 T cells. Immunity 29, 795–806 (2008).

63. Itano, A.A. et al. Distinct dendritic cell populations sequentially present antigen to CD4 T cells and stimulate different aspects of cell-mediated immunity. Immunity 19, 47–57 (2003).

64. Foy, T.M. et al. In vivo CD40-gp39 interactions are essential for thymus-dependent humoral immunity. II. Prolonged suppression of the humoral immune response by an antibody to the ligand for CD40, gp39. J Exp Med 178, 1567–1575 (1993).

65. Noelle, R.J. et al. A 39-kDa protein on activated helper T cells binds CD40 and transduces the signal for cognate activation of B cells. Proc Natl Acad Sci U S A 89, 6550–6554 (1992).

66. Poholek, A.C. et al. In vivo regulation of Bcl6 and T follicular helper cell development. J Immunol 185, 313–326 (2010).

67. Ballesteros-Tato, A. et al. Interleukin-2 inhibits germinal center formation by limiting T follicular helper cell differentiation. Immunity 36, 847–856 (2012).

68. Johnston, R.J., Choi, Y.S., Diamond, J.A., Yang, J.A. & Crotty, S. STAT5 is a potent negative regulator of TFH cell differentiation. J Exp Med 209, 243–250 (2012).

69. Ray, J.P. et al. The Interleukin-2-mTORc1 Kinase Axis Defines the Signaling, Differentiation, and Metabolism of T Helper 1 and Follicular B Helper T Cells. Immunity 43, 690–702 (2015).

70. DiToro, D. et al. Differential IL-2 expression defines developmental fates of follicular versus nonfollicular helper T cells. Science 361 (2018).

71. Walker, J.A. & McKenzie, A.N.J. T(H)2 cell development and function. Nat Rev Immunol 18, 121–133 (2018).

72. van Panhuys, N. et al. In vivo studies fail to reveal a role for IL-4 or STAT6 signaling in Th2 lymphocyte differentiation. Proc Natl Acad Sci U S A 105, 12423–12428 (2008).

73. Gillis, S., Ferm, M.M., Ou, W. & Smith, K.A. T cell growth factor: parameters of production and a quantitative microassay for activity. J Immunol 120, 2027–2032 (1978).

74. Morgan, D.A., Ruscetti, F.W. & Gallo, R. Selective in vitro growth of T lymphocytes from normal human bone marrows. Science 193, 1007–1008 (1976).

75. Laurence, A. et al. Interleukin-2 signaling via STAT5 constrains T helper 17 cell generation. Immunity 26, 371–381 (2007).

76. Yang, X.P. et al. Opposing regulation of the locus encoding IL-17 through direct, reciprocal actions of STAT3 and STAT5. Nat Immunol 12, 247–254 (2011).

77. Cote-Sierra, J. et al. Interleukin 2 plays a central role in Th2 differentiation. Proc Natl Acad Sci U S A 101, 3880–3885 (2004).

78. Kagami, S. et al. Stat5a regulates T helper cell differentiation by several distinct mechanisms. Blood 97, 2358–2365 (2001).

79. Liao, W. et al. Priming for T helper type 2 differentiation by interleukin 2-mediated induction of interleukin 4 receptor alpha-chain expression. Nat Immunol 9, 1288–1296 (2008).

80. Zhu, J., Cote-Sierra, J., Guo, L. & Paul, W.E. Stat5 activation plays a critical role in Th2 differentiation. Immunity 19, 739–748 (2003).

81. Zheng, W. & Flavell, R.A. The transcription factor GATA-3 is necessary and sufficient for Th2 cytokine gene expression in CD4 T cells. Cell 89, 587–596 (1997).

82. Zhu, J. et al. Conditional deletion of Gata3 shows its essential function in T(H)1-T(H)2 responses. Nat Immunol 5, 1157–1165 (2004).

83. Shi, M., Lin, T.H., Appell, K.C. & Berg, L.J. Janus-kinase-3-dependent signals induce chromatin remodeling at the Ifng locus during T helper 1 cell differentiation. Immunity 28, 763–773 (2008).

84. Bachus, H. et al. IL-6 prevents Th2 cell polarization by promoting SOCS3-dependent suppression of IL-2 signaling. Cell Mol Immunol 20, 651–665 (2023).

85. Ashour, D., et al. IL-12 from endogenous cDC1, and not vaccine DC, is required for Th1 induction. JCI Insight 5 (2020).

86. Harpur, C.M. et al. Classical Type 1 Dendritic Cells Dominate Priming of Th1 Responses to Herpes Simplex Virus Type 1 Skin Infection. J Immunol 202, 653–663 (2019).

87. Tatsumi, N. & Kumamoto, Y. Role of mouse dendritic cell subsets in priming naive CD4 T cells. Curr Opin Immunol 83, 102352 (2023).

